# A Budding Yeast Model and Screen to Define the Functional Consequences of Oncogenic Histone Missense Mutations

**DOI:** 10.1101/2021.10.02.462853

**Authors:** Laramie D. Lemon, Sneha Kannan, Kim Wai Mo, Miranda Adams, Haley G. Choi, Alexander O. D. Gulka, Elise S. Withers, Hasset T. Nurelegne, Valeria Gomez, Reina E. Ambrocio, Rhea Tumminkatti, Richard S. Lee, Morris Wan, Milo B. Fasken, Jennifer M. Spangle, Anita H. Corbett

## Abstract

Somatic missense mutations in histone genes turn these essential proteins into oncohistones, which can drive oncogenesis. Understanding how missense mutations alter histone function is challenging in mammals as mutations occur in a single histone gene. For example, described oncohistone mutations predominantly occur in the histone *H3.3* gene, despite the human genome encoding 15 H3 genes. To understand how oncogenic histone missense mutations alter histone function, we leveraged the budding yeast model, which contains only two H3 genes, to explore the functional consequences of oncohistones H3K36M, H3G34W, H3G34L, H3G34R, and H3G34V. Analysis of cells that express each of these variants as the sole copy of H3 reveals that H3K36 mutants show different drug sensitivities compared to H3G34 mutants. This finding suggests that changes to proximal amino acids in the H3 N-terminal tail alter distinct biological pathways. We exploited the caffeine sensitive growth of H3K36 mutant cells to perform a high copy suppressor screen. This screen identified genes linked to histone function and transcriptional regulation, including Esa1, a histone H4/H2A acetyltransferase; Tos4, a forkhead-associated domain-containing gene expression regulator; Pho92, an N6-methyladenosine RNA binding protein and Sgv1/Bur1, a cyclin-dependent kinase. We show that the Esa1 lysine acetyltransferase activity is critical for suppression of the caffeine sensitive growth of H3K36R mutant cells while the previously characterized binding interactions of Tos4 and Pho92 are not required for suppression. This screen identifies pathways that could be altered by oncohistone mutations and highlights the value of yeast genetics to identify pathways altered by such mutations.

## INTRODUCTION

Virtually every somatic cell within a eukaryotic organism contains identical genetic information; however, this identical information produces a plethora of cells with different morphologies and functions. Precise regulation of gene expression enables cells to have specific functions, structures, and biological responses. To enable dynamic responses via gene expression, DNA is packaged with histone proteins to form the basic chromatin unit of the nucleosome, which is a complex containing 147 base pairs of DNA wrapped around a histone octamer containing two copies each of the histones H2A, H2B, H3, and H4. DNA and histone proteins are modified by post-translational modifications (PTMs), which modulate DNA accessibility and regulate gene expression (1). Histones themselves are key regulators of nucleosome accessibility as a result of their dynamic acetylation, phosphorylation, and methylation. The exact combination, genome-wide localization, and dynamic addition or removal of histone PTMs contribute to the plasticity of gene expression, which is often deregulated in disease.

While the mechanisms by which characterized mutations in histone genes impact gene expression vary, expression of several disease-associated histone missense protein variants leads to chromatin remodeling and aberrant gene expression. To date, mutations in histone H3 have been most robustly characterized (2). In particular, data suggest that the H3K27M mutation decreases genome-wide repressive H3K27 methylation via impaired function of the polycomb repressive complex 2 (PRC2) (3–5). Loss of H3K27 methylation is visible across all genomic elements examined, including promoters, introns, intergenic regions, and 5’-and 3’-untranslated regions (UTRs) (3). Tissue characterized by H3K27M mutation also exhibits global DNA hypomethylation, leading to reduced promoter H3K27 methylation. Similarly, the H3K36M mutation is associated with a global reduction in H3K36 di-and tri-methylation, modulating gene expression and ultimately deregulating biological processes including cellular differentiation (6). Purified H3K36M-containing nucleosomes inhibit the activity of the H3K36-directed methyltransferases SETD2 and NSD2, and knockdown of these H3K36 methyltransferases phenocopies the effect H3K36M has on the epigenome and gene expression (7). These data suggest that while known histone mutations deregulate the epigenome and have drastic effects on gene expression, mechanisms by which histone mutations alter cell/biological function likely vary widely and these mutations have the potential to provide insight into specific histone functions.

The human genome encodes more than a dozen of each histone gene, H1, H2A, H2B, H3, and H4, which have arisen evolutionarily at least in part via gene duplication (8). All known and characterized disease-associated histone mutations (e.g., H3K27M, H3G34R, H3G34V, and H3K36M) act as dominant mutations, as a mutation in a single copy of one of the 15 encoded human H3 genes is sufficient to impart biological changes that impact cell growth (9). Oncohistone mutations that occur within N-terminal histone tails typically function through the deregulation of chromatin modification; in cases such as H3K27M or H3K36M histone variants, these mutations directly impact the ability of H3K27 or H3K36 to support methylation. As described above for H3K27M, the impact on histone post-translational modification occurs both in *cis* and *trans* (7); similar results are reported for H3K36M-driven cancers (10). These global changes to chromatin have the potential to deregulate transcriptional competence and subsequent gene expression on a large scale, which presents a new obstacle towards therapeutic development – how does one identify a deregulated transcriptional target(s) that is of consequence to cancer development? While recent studies have defined the dopamine receptor DRD2 as a therapeutically actionable gene target deregulated via H3K27M oncohistone expression (11), the identification of biologically meaningful cellular pathways and transcriptional changes that oncohistone-driven tumors are acutely dependent upon remains a pressing unmet clinical need.

While genomic alterations that support oncogenic growth have been identified in histone H1 (12) and histone H2B genes (13,14), most studies have focused on the histone H3 gene (9), in which typically one mutation is identified amongst the 15 genes, or 30 alleles, encoding H3 in the human genome. Oncogenic H3 mutations usually cluster in either of two genes encoding the H3 variant H3.3, which is the most evolutionarily conserved histone H3 variant (9). Studies on histones in model organisms including the budding yeast *S. cerevisiae* (15,16) and fission yeast *S. pombe* (17,18) can circumvent the complexity of studying the large number of histone gene copies in the human genome and provide insight into how histone mutations impart biologically relevant functional consequences. Such studies are informative as histones are highly conserved across species. Importantly, human and yeast H3 proteins have >97% identity (19). While the fission yeast *S. pombe* genome harbors three histone H3 genes, the budding yeast *S. cerevisiae* genome contains only two histone H3 genes, *HHT1* and *HHT2*, rendering either yeast model system amenable to the study of how the oncohistone changes in histone H3 alter histone function.

Previous studies in *S. pombe* have demonstrated that the oncogenic H3G34 mutations - H3G34V, H3G34W, and H3G34R - function through divergent mechanisms. The H3G34V mutation reduces H3K36 trimethylation (H3K36me3) (18), whereas the H3G34R mutation impairs Gcn5-mediated H3K36 acetylation (17). Experiments in *S. pombe* also highlight the different mechanisms of action of oncohistone mutations, as H3G34V-mutant yeast are sensitive to DNA-damaging agents, despite having intact homologous recombination-based DNA repair but are not sensitive to replicative stress (17). In contrast, H3G34R-mutant yeast are vulnerable to replicative stress but are not competent for homologous recombination DNA repair (18). These mutant oncohistones drive differential gene expression changes in yeast, which likely contribute to the contrasting sensitivities of the cells to stress. Such yeast studies highlight the concept that cancers that arise as a result of a specific oncohistone mutation are likely to benefit from different therapeutic modalities.

Similar results in yeast have been observed for various H3K36 amino acid substitutions. Expression of an H3K36R variant, which although structurally similar to the lysine normally present in this position, cannot be post-translationally modified (20), in *S. cerevisiae* renders yeast acutely sensitive to caffeine (21). Notably, H3K36R-expressing yeast cells exhibit a loss of H3K36 trimethylation (H3K36me3) and compromised growth in response to caffeine and rapamycin, which both inhibit growth factor and nutrient sensing pathways (21). The H3K36R mutation also alters alternative polyadenylation and pre-mRNA splicing in budding yeast (22,23). The H3K36R missense mutation is not linked to cancer, but it does alter the cellular transcriptional program and impair post-transcriptional processing in a *Drosophila melanogaster* model (20). In addition, an H3K36A mutation increases antisense transcription in budding yeast (24). All of these studies demonstrate the utility of employing budding yeast to explore the functional consequences of changing key amino acids within the histone H3 N-terminal tail.

Here we leverage the budding yeast model system to further elucidate the fundamental biological differences and vulnerabilities amongst established H3G34 and H3K36 oncohistones. Using a high copy suppressor screen approach to identify suppressors of oncohistone mutant growth phenotypes, we identify several genes linked to histone function, including the histone acetyltransferase, Esa1, the cyclin-dependent kinase, Sgv1/Bur1, a histone deacetylase complex-interacting protein, Tos4, and, finally, a protein that regulates mRNA stability and binds m^6^A RNA, Pho92. Notably, we find that the histone acetyltransferase activity of Esa1 is required for suppression of the caffeine sensitive growth of H3K36R mutant cells. Such approaches have the potential to guide novel mechanistic studies of human oncogenic histone mutations and suggest opportunities for therapeutic intervention for patients with tumors characterized by these oncohistones.

## MATERIALS AND METHODS

### Chemicals and media

All chemicals were obtained from Sigma-Aldrich (St. Louis, MO), United States Biological (Swampscott, MA), or Fisher Scientific (Pittsburgh, PA) unless otherwise noted. All media were prepared by standard procedures (25).

### S. cerevisiae strains and plasmids

All DNA manipulations were performed according to standard procedures (26). *S. cerevisiae* strains and plasmids used in this study are listed in Table S1. The PCR- and homologous recombination-based system for generating targeted mutations in histone genes in budding yeast cells has been described (27). Strains to model oncohistones—*hht2-K36R* (ACY2816), *hht2-K36M* (ACY2830), *hht2-G34W* (ACY2823), *hht2-G34L* (ACY2831), *hht2-G34R* (ACY2838), and *hht2-G34V* (ACY2841), which harbor mutations at the codons encoding the 36th or 34th histone H3 residue at the endogenous *HHT2* gene—were generated using the parental *hht2*Δ::*URA3* strain (yAAD165) and the strategy detailed previously (27,28). The endogenous *HHT1* gene in these oncohistone model strains was subsequently deleted and replaced via homologous recombination with a *kan*MX marker cassette to generate *hht2-K36R/M hht1*Δ (ACY2821, ACY2822) and *hht2-G34W/L/R/V hht1*Δ (ACY2825, ACY2833, ACY2840, ACY2846) strains. The *hht1*Δ (ACY2818) and *set2*Δ strain (ACY2851) were generated by deletion and replacement of the *HHT1* and *SET2* gene, respectively, via homologous recombination with a *kan*MX marker cassette. The YEp352 plasmids containing cloned *S. cerevisiae* genes identified in the high copy suppressor screen – *HHF2* (pAC4199), *HHT2* (pAC4201), *ESA1* (pAC4190), *TOS4* (pAC4196), *PHO92* (pAC4193), *SGV1* (pAC4187) – and *HHT1* (pAC4200) were generated by PCR amplification of each gene (5’ sequence/promoter, CDS, 3’UTR) from wildtype BY4741 genomic DNA with gene specific oligonucleotides (Integrated DNA Technologies) and conventional cloning via *Sal*I/*Sac*I or *Xho*I/*Sph*I (*TOS4*) into YEp352. All enzymes for PCR and cloning were obtained from New England BioLabs. The YEp352 plasmids containing *esa1/tos4/pho92/sgv1* variants with missense mutations in the Esa1 catalytic domain – *esa1-C304S* (pAC4191) and *esa1-E338Q* (pAC4192), the Tos4 Forkhead-associated domain –*tos4-N122A-N161A* (pAC4205), the Pho92 YTH domain – *pho92-W177A* (pAC4194) and *pho92-W231A* (pAC4195), and the Sgv1 catalytic domain – *sgv1-E107Q* (pAC4188), *sgv1-D213A* (pAC4189), *SUP3-E107Q* (pAC4212), and *SUP3-D213A* (pAC4213) were created using oligonucleotides containing the desired mutations (Integrated DNA Technologies), plasmid template - *ESA1* (pAC4190), *TOS4* (pAC4196), *PHO92* (pAC4193), SGV1 (pAC4187) or *SUP3* (pAC4132), and the QuikChange II Site-Directed Mutagenesis Kit (Agilent). The YEp352 plasmids containing *sgv1*Δ*2+8aa* (pAC4208), *SGV1-Myc* (pAC4209), *sgv1-E107Q-Myc* (pAC4210), and *sgv1-D213A-Myc* (pAC4211) were generated by PCR amplification of SGV1 5’sequence/CDS and 3’UTR products from *SGV1* plasmid template – *SGV1* (pAC4187), *sgv1-E107Q* (pAC4188), or *sgv1-D213A* (pAC4189) using oligonucleotides containing desired changes (Δ2+8aa; C-terminal Myc tag) and overlapping ends (Integrated DNA Technologies), and cloning into YEp352 linearized with *Eco*RI/*Hind*III using the NEBuilder Hifi DNA assembly cloning kit (New England BioLabs). All clones generated were sequenced to ensure that all cloned wildtype *S. cerevisiae* genes contained no mutations and all gene variants contained only the desired mutations.

### High copy suppressor screen

High-copy suppressors of the caffeine-sensitive growth of *hht2-K36R hht1*Δ cells (ACY2821) or *hht2-K36M hht1*Δ cells (ACY2822) were identified by transforming these cells with a YEp352-based high-copy *S. cerevisiae* genomic library (29) which was created by performing a *Sau*3AI partial digest of genomic *S. cerevisiae* DNA and cloning into the *Bam*HI site of the 2µm plasmid YEp352 (30). Transformants were grown on synthetic medium lacking uracil in the presence of 15 mM caffeine at 30°C for 5-10 days to select for those plasmids able to complement the caffeine-sensitive growth phenotype. Suppression by each plasmid (plasmid linkage) was confirmed by rescuing each suppressor plasmid and then retransforming them into the original H3K36 mutant to test for suppression of the slow growth on plates containing 15 mM caffeine.

### S. cerevisiae growth assays

To examine the growth of oncohistone model strains, wildtype (yAAD1253), *hht2-K36R/M* Δ*hht1* (ACY2821; ACY2822), *hht2-G34W/L/R/V hht1*Δ (ACY2825; ACY2833; ACY2840; ACY2846) and *hht1*Δ (ACY2818) strains were grown overnight at 30°C to saturation in 2 mL YEPD (yeast extract, peptone, dextrose) media. Cells were normalized to OD_600_ = 5, serially diluted in 10-fold dilutions, spotted on control YEPD media plates or YEPD media plates containing 15 mM caffeine or 3% formamide, and grown at 30°C and 18°C for 2-5 days. To test the effect of the genomic suppressor plasmids on the growth of oncohistone model strains and *set2*Δ cells in the presence of caffeine, wildtype (yAAD1253) cells transformed with YEp352 plasmid and *hht2-K36M hht1*Δ (ACY2822), *hht2-K36R hht1*Δ (ACY2821), and *set2*Δ (ACY2851) cells transformed with Vector (YEp352), *SUP3* (pAC4132), *SUP54* (pAC4145), *SUP67* (pAC4149), *SUP68* (pAC4150), or *SUP99* (pAC4160) plasmid were grown overnight at 30°C to saturation in 2 mL Ura-media containing 2% glucose. Cells were normalized by OD_600_ and serially diluted as previously described, spotted onto control YEPD media plates or YEPD media plates containing 15 mM caffeine, and grown at 30°C for 2-5 days. To test the effect of cloned suppressor genes and gene variants on the growth of *hht2-K36R hht1*Δ cells in the presence of caffeine, wildtype (yAAD1253) cells containing Vector (YEp352) and *hht2-K36R hht1*Δ (ACY2821) cells containing *HHF2* (pAC4199), *HHT2* (pAC4201), *HHT1* (pAC4200), *ESA1* (pAC4190), *esa1-C304S* (pAC4191), *esa1-E338Q* (pAC4192), *TOS4* (pAC4196), *tos4-R122A-N161A* (pAC4205), *PHO92* (pAC4193), *pho92-W177A* (pAC4194), *pho92-W231A* (pAC4195), *SGV1* (pAC4187), *sgv1*-Δ*2+8aa* (pAC4208), *SGV1-Myc* (pAC4209), *sgv1-E107Q-Myc* (pAC4210), *sgv1-D213A-Myc* (pAC4211), *SUP3-E107Q* (pAC4212), or *SUP3-D213A* (pAC4213) plasmid were grown overnight at 30°C to saturation in 2 mL Ura-media containing 2% glucose. As controls and for comparison, *hht2-K36R hht1*Δ (ACY2821) cells containing genomic suppressor plasmids were similarly grown. Cells were normalized by OD_600_ and serially diluted as previously described, spotted onto control YEPD plates or YEPD media plates containing 15 mM caffeine, and grown at 30°C for 2-5 days. For liquid growth assays, the indicated genotypes cells were grown in YEPD to mid-log phase and diluted into fresh media. The OD_600_ was recorded every 20 min in an Epoch2 microplate reader (BioTek) to determine doubling time. For the results shown, each sample was performed in three independent biological replicates with three technical replicates for each biological sample.

### Histone immunoblotting

To analyze histone H3K36me3 levels in oncohistone model strains, wildtype (yAAD1253), *hht2-K36R/M* Δ*hht1* (ACY2821; ACY2822), *hht2-G34W/L/R/V hht1*Δ (ACY2825; ACY2833; ACY2840; ACY2846), and *set2*Δ (ACY2851) strains were grown overnight at 30°C to saturation in 5 mL YEPD (yeast extract, peptone, dextrose) media. Cells were diluted in 50 mL YEPD to a starting OD_600_ = 0.1 and grown at 30°C to a final OD_600_ = 1.0. Cells were pelleted by centrifugation at 1,962 x *g* in 50 mL tubes and transferred to 2 mL screwcap tubes and pelleted by centrifugation at 16,200 x *g*. Pelleted cells were resuspended in 1 mL Lysis Buffer (10 mM Tris HCl, pH 8.0; 300 mM NaCl; 10% Glycerol; 0.1% IGEPAL® CA-630) supplemented with protease inhibitors [0.5 mM PMSF; Pierce™ Protease Inhibitors (Thermo Fisher Scientific)]. After addition of 500 µL acid washed glass beads, cells were disrupted in a Mini Bead Beater 16 Cell Disrupter (Biospec) for 3 × 30 sec at 25°C with 1 min on ice between repetitions. Cell debris was pelleted by centrifugation at 2,400 x *g* for 2 min at 4°C and protein lysate supernatant was transferred to fresh microfuge tube and clarified by centrifugation at 16,200 x *g* for 15 min at 4°C. The protein lysate was then transferred to a fresh tube. Protein lysate concentration was determined by Pierce BCA Protein Assay Kit (Life Technologies). Protein lysate samples (60 µg) in reducing sample buffer (50 mM Tris HCl, pH 6.8; 100 mM DTT; 2% SDS; 0.1% Bromophenol Blue; 10% Glycerol) were resolved on 4–20% Criterion™ TGX Stain-Free™ precast polyacrylamide gels (Bio-Rad). Protein lysate samples were transferred to nitrocellulose membranes (Bio-Rad) in Dunn Carbonate Buffer (10 mM NaHCO_3_, 3mM Na_2_CO_3_, pH 9.9, 20% methanol) at 22V for 90 min at room temperature. Histone H3K36me3 mark was detected with anti-H3K36me3 rabbit polyclonal antibody (ab9050; 1:1000; Abcam) and total histone H3 was detected with anti-H3 rabbit polyclonal antibody (ab1791; 1:5000; Abcam). Primary H3 rabbit antibodies were detected with secondary peroxidase-conjugated goat anti-rabbit IgG (111-035-003; 1:3000; Jackson ImmunoResearch Labs, Inc), ECL reagent, and ChemiDoc MP Imaging System (Bio-Rad).

### Quantitation of histone immunoblotting

The protein band intensities from immunoblots were quantitated using Image Lab software (Bio-Rad) and mean fold changes in protein levels were calculated in Microsoft Excel (Microsoft Corporation). The mean fold changes in histone H3K36me3 levels in oncohistone mutant cells relative to the wildtype control was calculated from two immunoblots. H3K36me3 band intensity was first normalized to total histone H3 band intensity and then normalized to H3K36me3 intensity in wildtype cells. The mean fold changes in H3K36me3 levels in oncohistone mutant cells relative to the wildtype control were graphed in GraphPad Prism 8 (GraphPad Software, LLC) with standard error of the mean error bars.

## RESULTS

### Missense mutations that model changes present in histone H3 confer different growth phenotypes

The N-terminal tails of the histone H3 protein are evolutionarily conserved with only a single conservative amino acid substitution (T->S) within the first 38 amino acids of the budding yeast compared to the human protein (Figure 1A). For this reason, the budding yeast system is valuable to model missense mutations that convert histone proteins into oncohistones. We focused on a set of missense mutations that alter K36 (K36M) or neighboring G34 (G34W, G34L, G34R, G34V), which are altered in various types of cancers, and an additional missense mutation that alters K36 and is associated with post-transcriptional regulation (21) (K36R).

**Figure 1.**
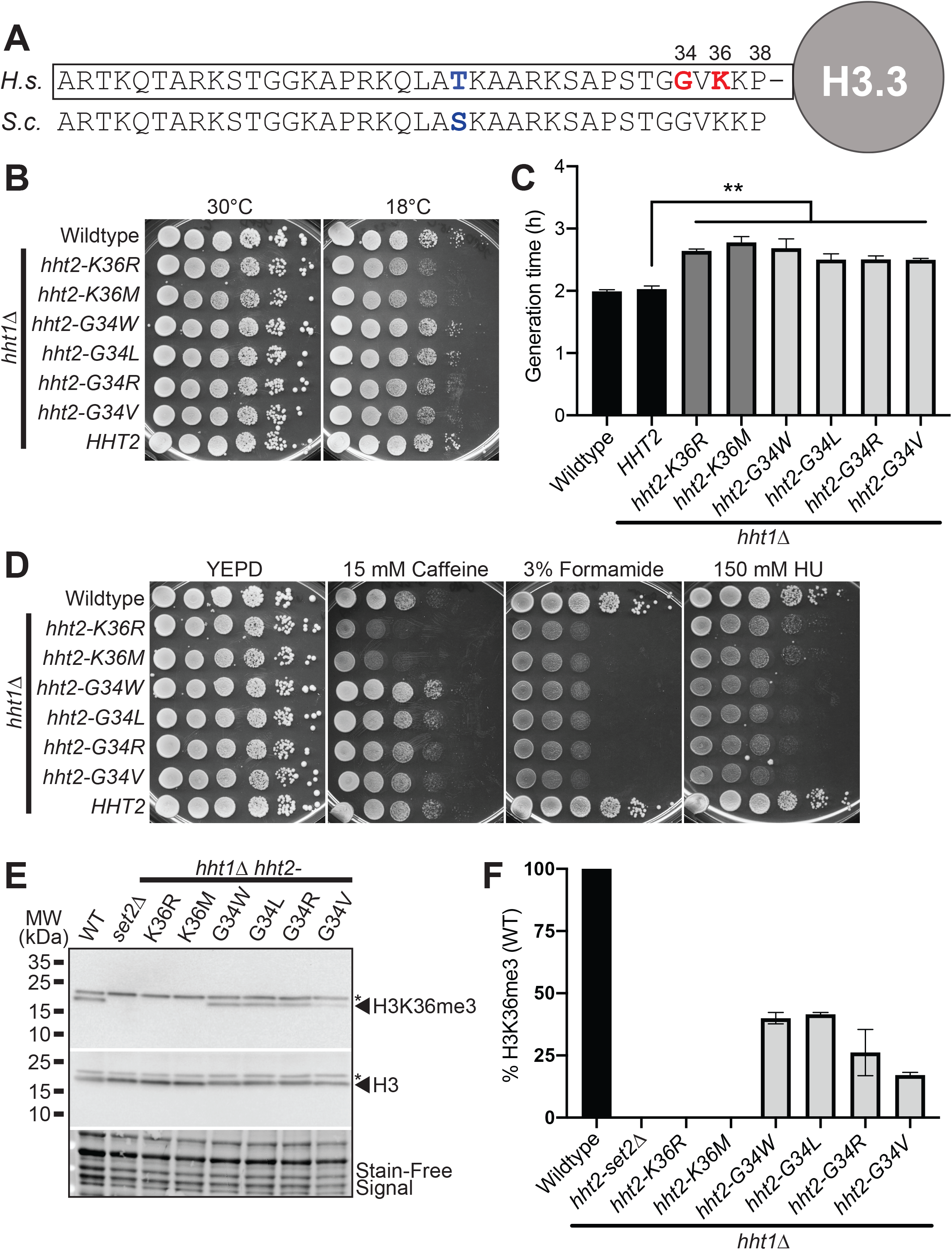
H3K36 and H3G34 oncohistone missense mutations within the conserved N-terminal tail of histone H3 cause diverse growth phenotypes in budding yeast. (A) The N-terminal tails of histone proteins are highly conserved as illustrated in the alignment shown for human (*H. sapiens*, *H.s.*) histone H3.3 and *S. cerevisiae* (*S.c.*) Hht1/Hht2. The positions of K36 and G34, which are the residues altered in the oncohistones modeled here are indicated by the red and the single conservative amino acid change from threonine (T) in human to serine (S) in budding yeast at position 22 is indicated in blue. (B) The H3K36 histone mutant cells, but not H3G34 mutant cells, show cold-sensitive growth. The H3K36R/M mutant cells (*hht2-K36R/M hht1*Δ) show impaired growth at 18°C compared to wildtype cells. The H3K36 and H3G34 mutant cells express each histone variant as the sole copy of histone H3. Wildtype, H3K36 mutant cells (*hht2-K36R/M hht1*Δ), H3G34 mutant cells (*hht2-G34W/L/R/V hht1*Δ), and *HHT2 hht1*Δ cells were serially diluted and spotted on YEPD media plates and grown at the permissive temperature of 30°C and cold temperature of 18°C. (C) A growth curve was employed to analyze growth rate (Figure S1). Doubling times were obtained for control Wildtype and *HHT2 hht1*Δ cells (black), H3K36 mutant cells (*hht2-K36R/M hht1*Δ) (gray), and H3G34 mutant cells (*hht2-G34W/L/R/V hht1*Δ) (light gray). The doubling time was increased for each of the H3 mutants analyzed compared to either control. Statistical significance (** p<0.001) was determined by using a one-tailed Student’s T-test. (D) The H3K36 histone mutant cells show caffeine sensitive growth, whereas H3G34 mutants cells show hydroxyurea (HU) sensitive growth. The H3K36R/M mutant cells (*hht2-K36R/M hht1*Δ) show impaired growth on caffeine plates, whereas the H3G34 mutant cells (*hht2-G34W/L/R/V hht1*Δ) show impaired growth on hydroxyurea (HU) plates compared to wildtype cells. All H3K36 and H3G34 mutant cells analyzed show severely impaired growth on formamide plates. Wildtype, H3K36 mutant cells (*hht2-K36R/M hht1*Δ), H3G34 mutant cells (*hht2-G34W/L/R/V hht1*Δ), and *HHT2 hht1*Δ cells were serially diluted and spotted on a control YEPD media plate and YEPD plates containing 15 mM Caffeine, 3% Formamide, or 150 mM Hydroxyurea (HU). and grown at 30°C. (E,F) Immunoblotting was performed to analyze levels of H3K36 trimethylation (H3K36me3) in budding yeast oncohistone models. Wildtype, control *set2*Δ, H3K36 mutant cells (*hht2-K36R/M hht1*Δ), and H3G34 mutant cells (*hht2-G34W/L/R/V hht1*Δ) were grown and analyzed as described in Materials and Methods to detect H3K36me3 and total H3. Stain-Free Signal is included as a total loading control. (E) A representative immunoblot demonstrating loss of H3K36me3 in the control *set2*Δ and H3K36 mutant cells (*hht2-K36R/M hht1*Δ) with a decrease in H3K36me3 in the H3G34 mutant cells. A non-specific band detected by the antibody is indicated by the asterisk. Molecular weight markers are indicated to the left. (F) A bar graph presents the results from two independent immunoblot experiments as shown in (E). The level of H3K36me3 was normalized to Stain-Free Signal and total H3 levels. The level of H3K36me3 detected in control WT cells was set to 100% and all other samples were calculated relative to this level of H3K36me3.

We created *S. cerevisiae* H3 mutant models where each histone variant is expressed as the sole copy of histone H3 and then assessed cell growth using a serial dilution assay. As shown in Figure 1B, in an end point solid media growth assay, all histone mutants show growth comparable to either control wildtype (WT) cells or control cells lacking the *HHT1* gene (*HHT2 hht1*Δ), which is absent in the oncohistone mutant models that contain *HHT2* as the sole histone H3 gene. At cold temperature (18°C), both the H3K36R and H3K36M mutant cells show a modest growth defect (Figure 1B) with no change in growth detected for any of the H3G34 variants. To extend this analysis, we performed a liquid growth assay, which captures changes in the rate of growth. At the permissive temperature of 30°C, differences in the growth rate for each of the oncohistone mutants relative to WT control cells are illustrated by increased doubling time (Figure 1C) and slower growth rate (Figure S1). These data demonstrate that these oncohistone mutants exhibit changes in cell growth.

To explore whether the oncohistone models exhibit other changes, we then tested for growth defects when cells are grown on media containing chemicals that disrupt different cellular pathways (Figure 1D). Caffeine impairs cellular stress response/TOR signaling (31); formamide alters RNA metabolism (32), and hydroxyurea (HU) impairs DNA synthesis (33). Results of this analysis reveal that amino acid substitutions at H3K36 cause sensitivity to growth on media containing caffeine, which is consistent with previous results showing that loss of the H3K36 methyltransferase Set2 (34) and expression of the H3K36R variant confer sensitivity to caffeine (21). All of the histone H3 mutants are sensitive to formamide, while the H3G34 mutants are more sensitive to hydroxyurea than the H3K36 mutants.

Analysis of H3K36 trimethylation (H3K36me3) in the oncohistone mutant cells reveals that H3K36R and H3K36M mutants show no detectable H3K36me3, as expected and similar to control *set2*Δ cells, whereas H3G34 mutants show decreased H3K36me3 relative to wildtype cells (Figure 1E, 1F). Notably, amongst the H3G34 mutants, H3G34V shows the greatest decrease in H3K36me3. These results suggest that the drug sensitive growth of the oncohistone mutants is linked to an altered epigenome. In addition, these data reveal that different changes to residues within the N-terminal tail of histone H3 confer distinct growth defects.

### A high copy suppressor screen to identify functional links to missense mutations present in oncohistones

To explore the molecular basis of the growth defects conferred by missense mutations in the histone H3 gene, we performed a high copy suppressor screen. As described in Materials and Methods, either H3K36M or H3K36R mutant cells were transformed with a high copy genomic library (29) and plated on media containing 15 mM caffeine. Suppressors that enhanced growth of these mutants in the presence of caffeine were isolated and selected for further analysis. Figure 2A shows results for a set of suppressor clones (*SUP*) identified in the screen that suppress the caffeine sensitive growth of the H3K36M oncohistone model. While the screen was not saturated, we did identify several clones multiple times. We validated each suppressor by plasmid rescue and retransformation.

**Figure 2.**
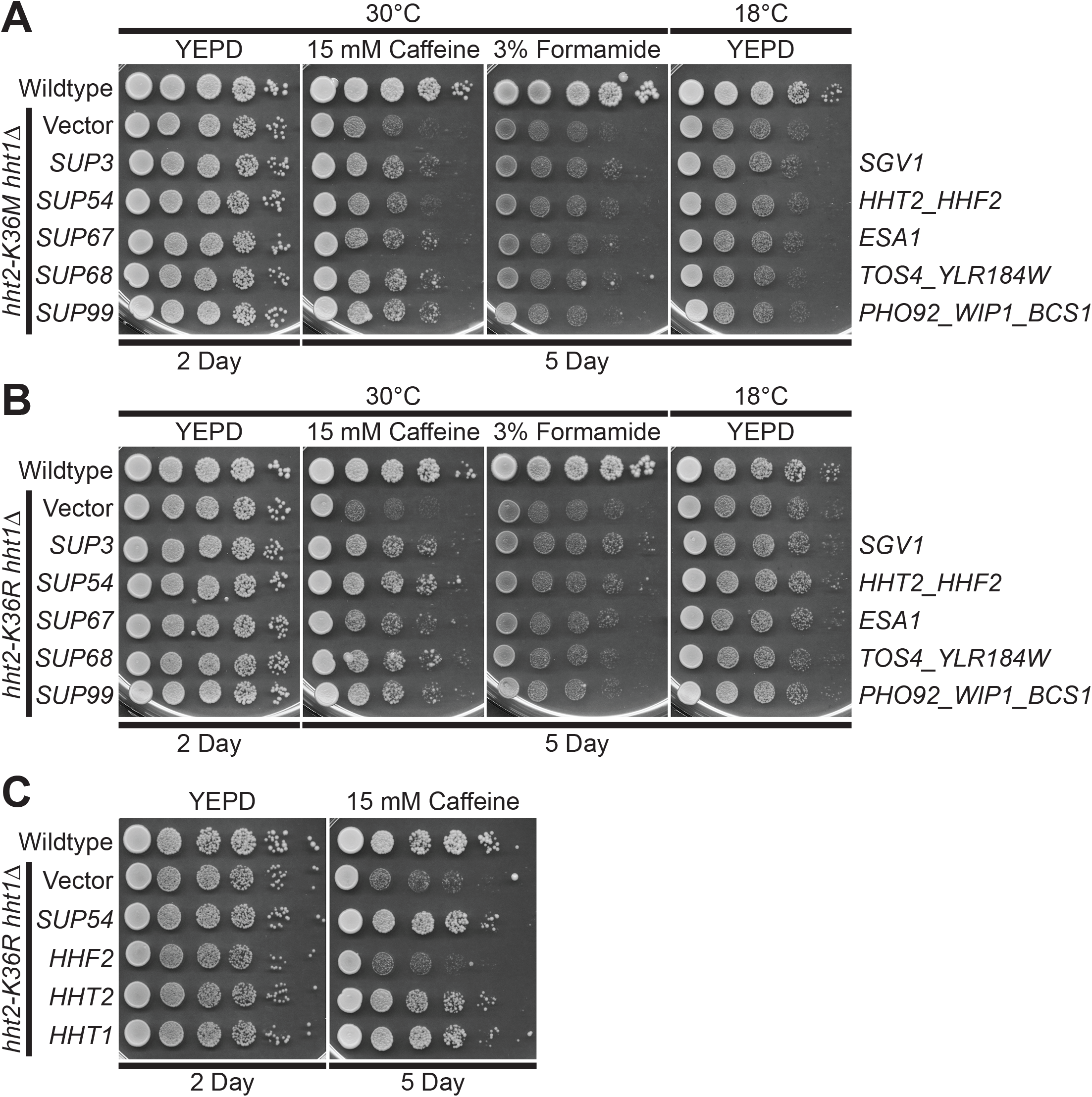
A high copy suppressor screen for suppression of the caffeine sensitive growth of H3K36 histone mutant cells identifies five suppressors. As described in Materials and Methods, a high copy suppressor screen was performed to identify genes that suppress the caffeine sensitive growth of H3K36M and/or H3K36R mutant cells. (A, B) The five genomic suppressor (*SUP*) plasmids identified from the screen suppress the caffeine sensitive growth of (A) H3K36M and (B) H3K36R histone mutant cells. The H3K36M/R mutant cells (*hht2-K36M/R hht1*Δ) containing *SUP3/54/67/68/99* suppressor plasmids show improved growth on caffeine plates compared to cells containing vector alone. The H3K36M/R cells containing suppressor plasmids do not show any effect on growth at 18°C and little effect on formamide plates; however, cells containing the *SUP3* suppressor show slightly improved growth compared to cells containing vector alone on formamide plates. Sequencing of the genomic suppressor plasmids revealed the identity of the gene(s) encoded on the clones: *SUP3* (*SGV1*), *SUP54* (*HHT2, HHF2*), *SUP67* (*ESA1*), *SUP68* (*TOS4, YLR184W*), and *SUP99* (*PHO92, WIP1, BCS1*) –indicated to right. Wildtype cells transformed with vector and H3K36M/R mutant cells (*hht2-K36M/R hht2*Δ) transformed with vector, *SUP3*, *SUP54*, *SUP67*, *SUP68*, or *SUP99* plasmid were serially diluted and spotted onto YEPD plates and YEPD plates containing 15 mM Caffeine or 3% Formamide and grown at indicated temperatures. (C) The *HHT2* histone H3 gene, but not the *HHF2* histone H4 gene, suppresses the caffeine sensitive growth of H3K36R histone mutant cells. The H3K36R mutant cells (*hht2-K36R hht1*Δ) containing an *HHT2* plasmid show improved growth compared to cells with vector alone on a caffeine plate, similar to cells containing the *SUP54* suppressor, which harbors both *HHT2* and *HHF2*. The H3K36R cells containing an *HHF2* plasmid do not show improved growth on a caffeine plate. The H3K36R cells containing an *HHT1* plasmid, containing the other *<u>S. cerevisiae</u>* histone H3 gene, show improved growth on a caffeine plate. Wildtype cells transformed with vector and H3K36R mutant cells (*hht2-K36R hht2*Δ) transformed with vector, *SUP54*, *HHF2, HHT2,* or *HHT1* plasmid were serially diluted and spotted onto a control YEPD plate and YEPD plate containing 15 mM Caffeine and grown at 30°C.

As expected in a suppressor screen, we identified a suppressor that contains one of the two budding yeast H3 genes, *HHT2*. Beyond *HHT2*, which validates the screen, we focused our analysis on four suppressor clones that illustrate the power of this approach to define the biological pathways altered in oncohistone model cells. These suppressors encode the cyclin-dependent protein kinase, Sgv1/Bur1 (35,36), the catalytic subunit of the NuA4 histone acetyltransferase complex, Esa1 (37), the gene expression regulator, Tos4 (38,39), which contains a forkhead-associated (FHA) domain that interacts with Rpd3L and Set3 histone deacetylase complexes (40), and a post-transcriptional regulator of phosphate and glucose metabolism, Pho92 (41) (Figure 2A,B).

To begin to assess whether the suppressors identified are linked to post-translational modification of H3K36, we tested whether the suppressors identified can also suppress a yeast mutant that expresses the conservative H3K36R variant as the sole copy of histone H3. As shown in Figure 2B, each of the suppressors identified also suppresses the caffeine sensitive growth of H3K36R cells. Thus, the suppressors identified for further analysis suppress the growth phenotypes of both H3K36M and H3K36R mutant cells in a similar manner.

As a first step to validate the screen and the suppressors identified, we focused on the suppressor clone containing the histone H3 gene *HHT2* (*SUP54*). This genomic clone contains one of the two budding yeast histone H3 genes (*HHT2*) and one of the two budding yeast histone H4 genes (*HHF2*). We subcloned the *HHT2* and *HHF2* genes and tested them independently for suppression of the caffeine sensitive growth of the H3K36R cells (Figure 2C). This analysis reveals that *HHT2* rescues the growth defect of these cells, while the *HHF2* clone confers no rescue as cells show growth comparable to the Vector alone control. We also confirmed that, as expected, the other budding yeast histone H3 gene, *HHT1*, rescues the caffeine sensitive growth of H3K36R mutant cells (Figure 2C).

The *S. cerevisiae ESA1* gene encodes an essential, evolutionarily conserved lysine acetyltransferase that acetylates lysine residues within the N-terminal tail of histone H4 as well as histone H2A (37,42). The human orthologue of Esa1 is the KAT5/TIP60 protein (43). As illustrated in the domain structure shown in Figure 3A, Esa1 is a member of the MYST (MOZ[KAT6A], <u>Y</u>BF2/Sas3, <u>S</u>as2, <u>T</u>IP60[KAT5]) family of lysine acetyl transferases (44,45). Esa1 also contains an N-terminal chromo domain (46). These same functional domains are conserved within the human KAT5/TIP60 protein (43).

**Figure 3.**
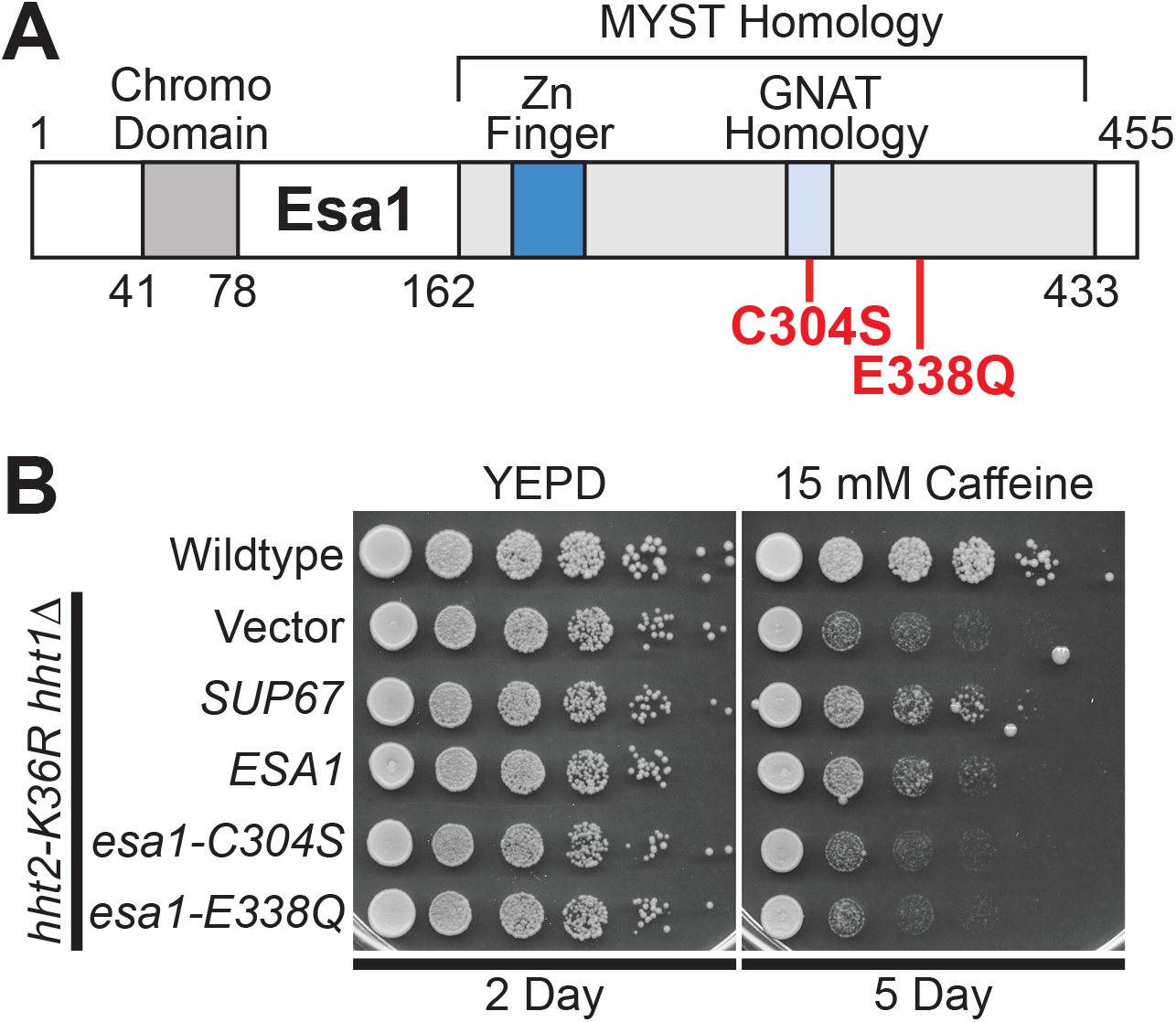
The lysine acetyltransferase *ESA1* suppresses the caffeine sensitive growth of H3K36R histone mutant cells and requires the catalytic activity of Esa1 for suppression. *SUP67* includes a single intact gene *ESA1*, which encodes a histone H4/H2A lysine acetyl transferase of the MYST (Moz, <u>Y</u>BF2, <u>S</u>as2, <u>T</u>ip) family. (A) The domain structure of Esa1 is shown. The MYST homology domain contains both a Zinc Finger (Zn Finger) domain and a Gcn5-related N-acetyltransferases (GNAT) Homology domain. Esa1 also contains an N-terminal Chromo Domain. The amino acid changes created to impair the lysine acetyltransferase activity of Esa1, C304S and E338Q, which are based on previous work (47) and located in the MYST homology domain, are shown in red. (B) *ESA1* suppresses the caffeine sensitive growth of H3K36R mutant cells similar to the high copy suppressor, *SUP67*, but catalytically inactive mutants of Esa1 do not suppress. The H3K36R cells (*hht2-K36R hht1*Δ) containing an *ESA1* plasmid show improved growth compared to cells with vector alone on a caffeine plate, similar to cells containing the *SUP67* suppressor, but cells containing the catalytically inactive mutant *esa1-C304S* or *esa1-E338Q* plasmid do not show improved growth. Wildtype cells transformed with vector and H3K36R mutant cells (*hht2-K36R hht2*Δ) transformed with vector, *SUP67*, *ESA1, esa1-C304S,* or *esa1-E338Q* plasmid were serially diluted and spotted onto a control YEPD plate and YEPD plate containing 15 mM Caffeine and grown at 30°C for indicated days.

We confirmed that overexpression of *ESA1*, the only intact gene present in the *SUP67* suppressor, can suppress the caffeine sensitive growth of H3K36R mutant cells. To assess whether the lysine acetyltransferase function of Esa1 is critical for this growth suppression, we took advantage of two previously characterized catalytic mutants (Figure 3A), *esa1-C304S* and *esa1-E338Q* (44,47). Each of these amino acid substitutions eliminates the acetyltransferase activity of Esa1 without a significant impact on steady-state protein level (47). As shown in Figure 3B, neither of these catalytic mutants of Esa1 can suppress the growth of H3K36R mutant cells. Thus, the acetyltransferase activity of Esa1 is required to suppress the growth defects associated with H3K36 mutants.

The Tos4 protein is a gene expression regulator that plays a role in gene expression homeostasis (38,39). As depicted in Figure 4A, Tos4 contains a forkhead-associated (FHA) domain, which mediates interactions with histone deacetylase (HDAC) complexes, Rpd3L and Set3 (40). We cloned *TOS4,* as the *SUP68* suppressor clone slightly truncates the Tos4 open reading frame and also contains an uncharacterized gene *YLR184W.* As shown in Figure 4B, overexpression of *TOS4* suppresses the caffeine sensitive growth of H3K36R mutant cells. To test whether a functional forkhead-associated (FHA) domain is required for Tos4-mediated growth suppression, we exploited amino acid substitutions R122A and N161A in the Tos4 FHA domain that disrupt Tos4 interaction with HDAC complexes, Rpd3L and Set3 (40). We created a Tos4 variant that contains both these amino acid substitutions (Figure 4A); however, this double *tos4-R122A-N161A* mutant still robustly suppresses growth defects of H3K36R mutant cells, suggesting that a HDAC independent, alternative function of Tos4 mediates this suppression.

**Figure 4.**
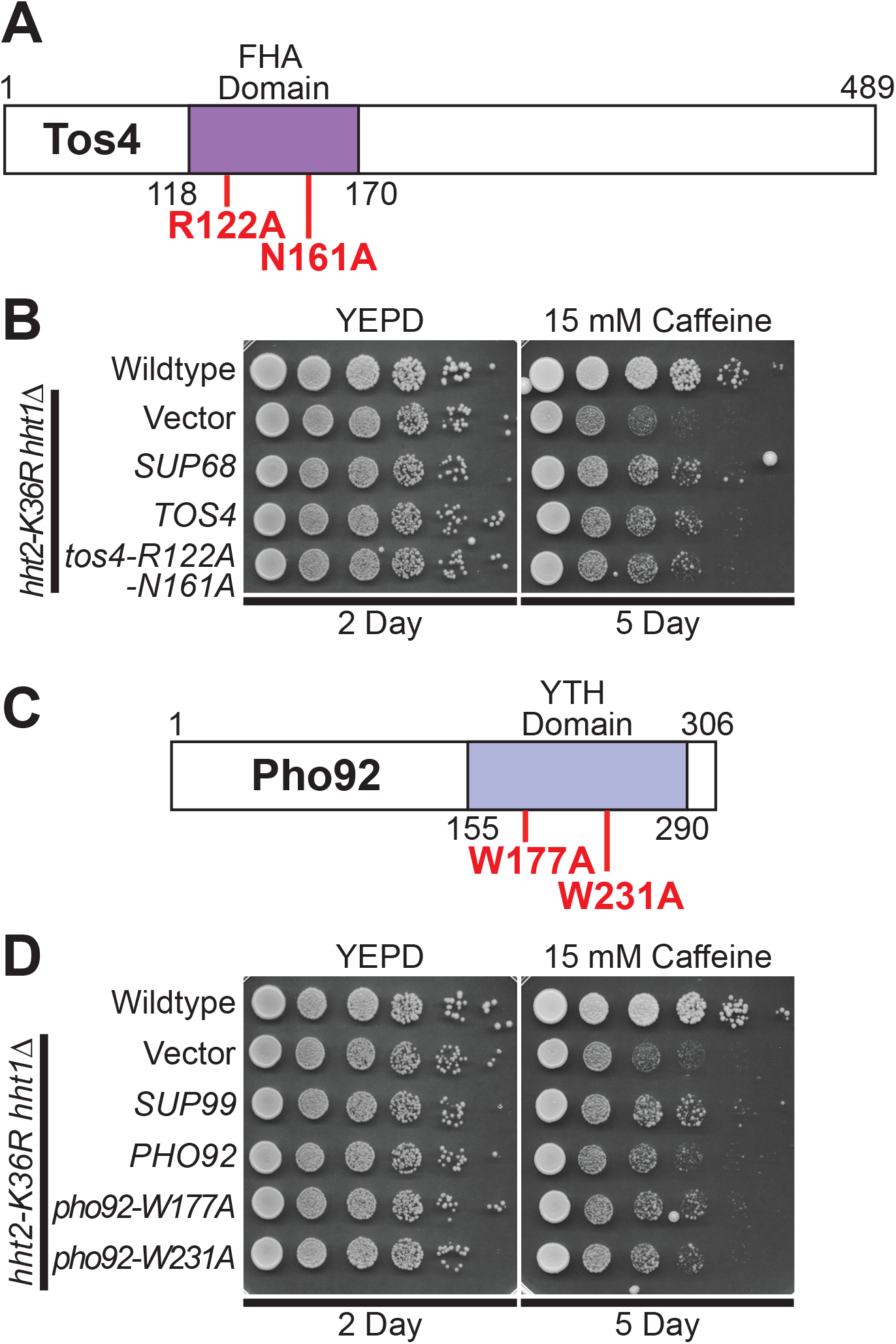
The gene expression regulator *TOS4* and m^6^A RNA binding protein *PHO92* suppress the caffeine sensitive growth of H3K36R histone mutant cells, but Tos4 interaction with HDAC complexes and Pho92 m^6^A RNA binding are not required for suppression. (A) The Tos4 protein is a gene expression regulator that contains a forkhead-associated (FHA) domain, which is a phosphopeptide recognition domain found in many regulatory proteins. The amino acids changes generated to disrupt the interaction with the histone deacetylase (HDAC) complexes, Rpd3L and Set3 (39,40), R122A and N161A, which are located in the FHA domain, are shown in red. (B) *TOS4* suppresses the caffeine sensitive growth of H3K36R mutant cells to the same extent as the high copy suppressor, *SUP68*, and a FHA domain double mutant of Tos4 that disrupts interaction with Rpd3L and Set3 HDACs remains competent to suppress the cells. The H3K36R cells (*hht2-K36R hht1*Δ) containing a *TOS4* plasmid show improved growth compared to cells with vector alone on a caffeine plate, similar to cells containing the *SUP68* suppressor, which contains uncharacterized open reading frame, *YLR184W*, in addition to *TOS4*. The H3K36R cells containing an FHA domain double mutant of Tos4, *tos4-R122A-N161A*, show improved growth compared to cells with vector alone on a caffeine plate, similar to cells containing *TOS4*. (C) The Pho92 protein contains a YT521-B homology (YTH) domain, which is an evolutionarily conserved m^6^A-dependent RNA binding domain (49). The amino acid changes made to impair the binding of Pho92 to m^6^A RNA, W177A and W231A, which alter key tryptophan residues in the m^6^A binding pocket (49) located in the YTH domain, are indicated in red. (D) *PHO92* suppresses the caffeine sensitive growth of H3K36R mutant cells similar to the high copy suppressor, *SUP99*, but the m^6^A RNA binding function of Pho92 is not required for suppression. The H3K36R cells (*hht2-K36R hht1*Δ) containing a *PHO92* plasmid show improved growth compared to cells with vector alone on a caffeine plate, similar to cells containing the *SUP99* suppressor, which contains *WIP1* and *BCS1* genes in addition to *PHO92*. The H3K36R cells containing the m^6^A RNA binding mutant *pho92-W177A* or *pho92-W231A* plasmid show improved growth compared to cells with vector alone on a caffeine plate, similar to cells containing *PHO92*. Wildtype cells transformed with vector and H3K36R mutant cells (*hht2-K36R hht2*Δ) transformed with vector, *SUP68*, *TOS4, tos4-R122A-N161A, SUP99*, *PHO92*, *pho92-W177A*, or *pho92-W231A* plasmid were serially diluted and spotted onto a control YEPD plate and YEPD plate containing 15 mM Caffeine and grown at 30°C for indicated days.

The Pho92 protein is a member of an evolutionarily conserved family of proteins that contains a C-terminal YTH domain (41) (Figure 4C). The YTH domain can recognize and bind m^6^A-containing RNA (48), serving as the primary “readers” of this post-transcriptional modification of RNA. The high copy suppressor containing *PHO92* (*SUP99*) also contained two other genes, *WIP1* and *BCS1*, so we cloned the *PHO92* gene and found that *PHO92* is indeed a high copy suppressor of the caffeine sensitive growth of H3K36R mutant cells (Figure 4D). To determine whether YTH-mediated interaction with m^6^A-containing RNA is required for this suppression, we altered conserved tryptophan residues that are critical to form the binding pocket for m^6^A (49,50), creating *pho92-W177A* and *pho92-W231A* (Figure 4C). As shown in Figure 4D, both of these *pho92* mutants still suppress the caffeine sensitive growth of the H3K36R cells, strongly suggesting that the YTH domain is not required for this suppression.

The Sgv1 protein (also termed Bur1) is an evolutionarily conserved cyclin-dependent kinase (35), which is closely related to human CDK9 (51,52), that is required for efficient transcription elongation by RNA polymerase II (RNAPII) (53). Importantly, Sgv1/Bur1 phosphorylates the Rbp1 linker region between the RNAPII body and C-terminal domain (CTD), facilitating recruitment of the elongation factor Spt6 (51), and phosphorylates the C-terminal region (CTR) of elongation factor Spt5 (54). Consistent with this function, the Sgv1 protein contains a protein kinase domain (Figure 5A). Although *SGV1* was the sole intact gene present on the *SUP3* suppressor, the C-terminal end of the Sgv1 open reading frame is truncated by two amino acids (loss of LY) with a predicted addition of 8 amino acids (PRVPSSNS) due to translation into the multiple cloning site of the vector (Figure 5A). Thus, we cloned *SGV1* and tested for suppression of the caffeine sensitive growth of H3K36R mutant cells. Surprisingly, the wildtype *SGV1* clone does not suppress this growth defect (Figure 5B). We then considered the possibility that the C-terminal loss of either two amino acids (Δ2) or addition of the eight amino acids (+8AA) could confer suppression through a dominant mechanism. We thus reengineered a clone to independently produce the same Sgv1 variant (Figure 5A, termed sgv1Δ2+8 amino acids or sgv1Δ2+8aa) as the one identified in the high copy suppressor screen. As shown in Figure 5B, this Sgv1 variant suppresses the H3K36R growth defect on caffeine. We then considered the possibility that any C-terminal extension of Sgv1 is sufficient to mediate this suppression, so we appended a Myc tag (EQKLISEEDL) to the C-terminus of the wildtype *SGV1* open reading frame (Figure 5A termed SGV1-Myc). Interestingly, this C-terminally Myc-tagged Sgv1 protein also suppresses the caffeine-sensitive growth of H3K36R mutant cells (Figure 5B). Importantly, all the engineered clones contain the endogenous *SGV1* 3’UTR so differences should not be due to changes in 3’-UTR-mediated regulation.

**Figure 5.**
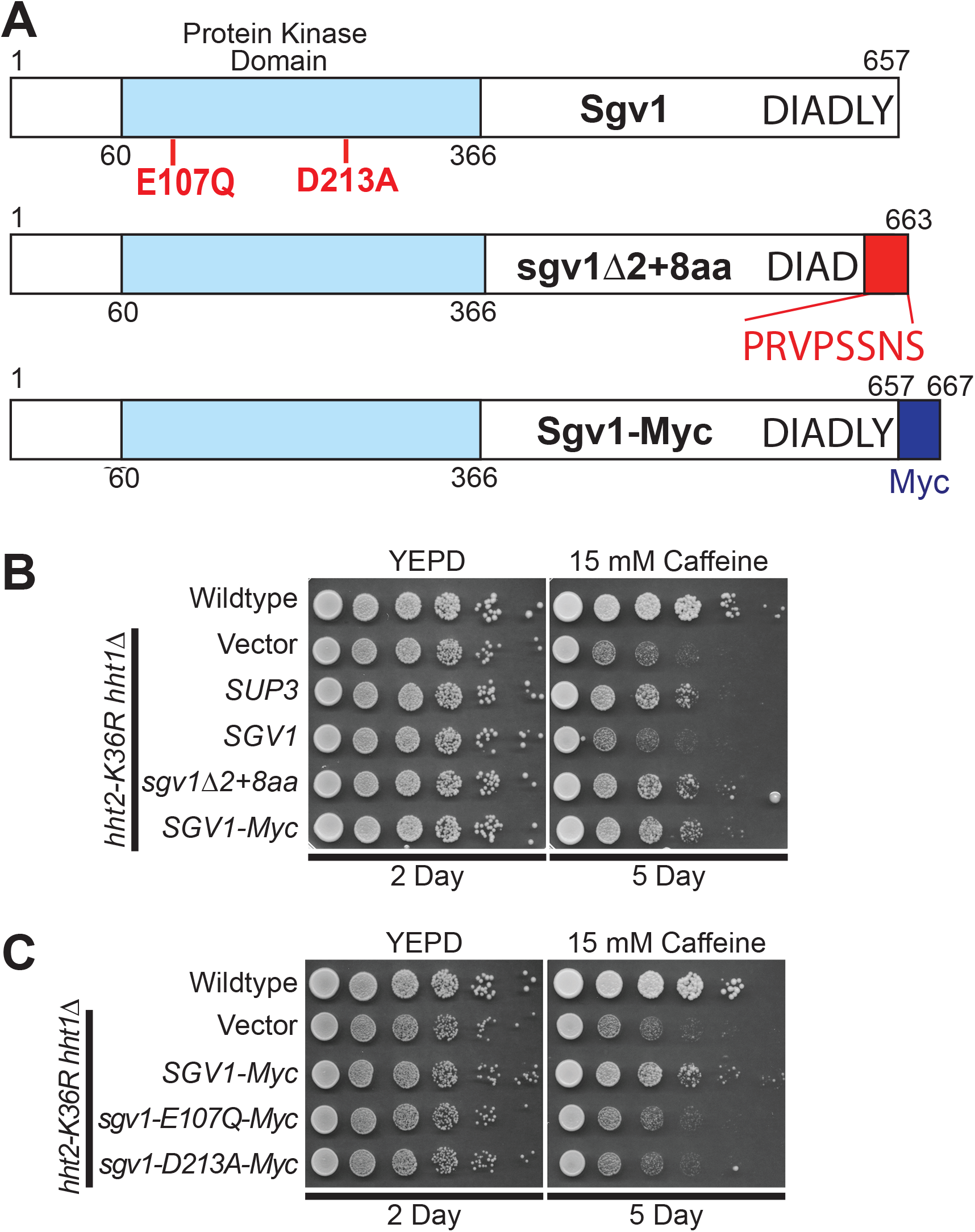
Wildtype *SGV1* does not suppress the caffeine sensitive growth of H3K36 histone mutants, but Sgv1 with a C-terminal extension can suppress. (A) The Sgv1/Bur1 kinase is a cyclin-dependent kinase that contains a protein kinase domain with homology to human CDK9 (52). The amino acid substitutions generated to inactivate the catalytic function of Sgv1/Bur1 (53), E107Q and D213A, which are located in the protein kinase domain, are shown in red. The genomic suppressor clone *SUP3* identified in the high copy suppressor screen encodes a slightly truncated Sgv1 protein, which lacks the last two amino acids (LY) and translates into the multiple cloning site of the vector (YEp352) to add an additional 8 amino acids (PRVPSSNS), which are indicated in red at the C-terminus on the domain structure shown and labeled sgv1Δ2+8aa. An additional Sgv1 variant was engineered to contain an in-frame C-terminal Myc tag (EQKLISEEDL), which is labeled Sgv1-Myc. (B) Unlike the high copy suppressor, *SUP3*, which suppresses the caffeine sensitive growth of H3K36R mutant cells, the wildtype *SGV1* clone does not suppress the cells; however, an *SGV1* clone containing the C-terminal addition of either the suppressor screen-associated changes (Δ2+8 amino acids), *sgv1*Δ*2+8aa*, or a Myc tag, *SGV1-Myc*, suppresses growth on caffeine. (C) While the control *SGV1-Myc*, suppresses the caffeine sensitive growth of H3K36 mutant cells, *SGV1-Myc* containing either the catalytic amino acid substitution E107Q or D213A (53) does not suppress the caffeine sensitive growth defect of H3K36R mutant cells.

Having found that addition of Myc tag to the C-terminus of Sgv1 creates a functional suppressor of the H3K36 growth defect on caffeine (Figure 5B), we next tested whether the catalytic activity of Sgv1 is required for this suppression. In this Sgv1-Myc variant, we generated two previously characterized mutants that impair the catalytic function of Sgv1/Bur1 – *sgv1-E107Q* and *sgv1-D213A* (53). As shown in Figure 5C, when either of these changes that impair the catalytic activity of Sgv1 are introduced into Sgv1-Myc, no growth suppression is observed, arguing that the catalytic function of Sgv1 is required to suppress the caffeine-sensitive growth of H3K36R mutant cells. In support of this conclusion, introduction of the E107Q or D213A catalytic mutations into the Sgv1-containing *SUP3* suppressor also impairs its ability to suppress H3K36R growth on caffeine (Figure S2). Thus, both the catalytic function and a change to the C-terminus of Sgv1 are required to suppress the growth of the oncohistone mutant cells.

Given that these suppressors were identified in a screen for genes that could suppress growth defects of either H3K36M or H3K36R mutant cells on plates containing caffeine, the mechanism of suppression is likely linked to post-translational modification of this critical Lys36 residue in histone H3. To test this idea, we examined whether the identified genomic suppressors can suppress the caffeine sensitive growth of budding yeast cells lacking the H3K36 histone methyltransferase, Set2 (21,34). Four of the five identified genomic suppressors - *SUP3* (*SGV1*), *SUP67* (*ESA1*), *SUP68* (*TOS4*), and *SUP99* (*PHO92*) - can suppress the caffeine sensitive growth of *set2*Δ cells. As expected suppressor *SUP54* (*HHT2*)does not suppress the *set2*Δ caffeine sensitive growth defect (Figure 6). This result is expected because an increase in the dose of histone H3 gene in the absence of a functional histone methyltransferase would not be expected to overcome the growth defect caused by loss of the methyltransferase and the accompanying methylation of H3K36. These data functionally link these suppressors to proper methylation of H3K36.

**Figure 6.**
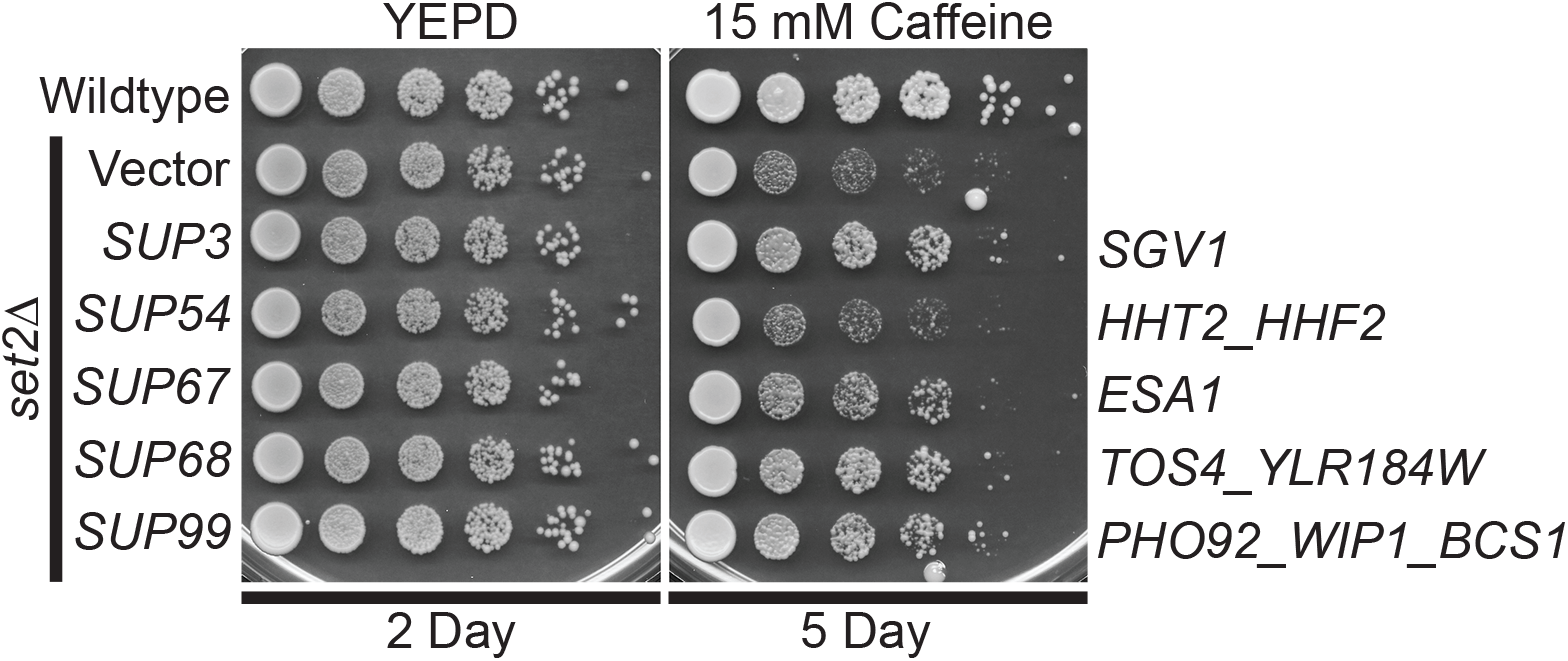
The identified genomic suppressors of H3K36R/M histone mutants, containing *SGV1*, *ESA1*, *TOS4*, and *PHO92*, suppress the caffeine sensitive growth of *set2*Δ H3K36 histone methyltransferase mutant cells. The *set2*Δ mutant cells containing *SUP3* (*SGV1*), SUP*67* (*ESA1*), *SUP68* (*TOS4*), and *SUP99* (*PHO92*) suppressor plasmids show improved growth on a caffeine plate compared to cells containing vector alone, but cells containing *SUP54* (*HHT2*) do not improve growth. The gene(s) encoded on the suppressor clones are indicated to right. Wildtype cells transformed with vector and *set2*Δ mutant cells transformed with vector, *SUP3*, *SUP54*, *SUP67*, *SUP68*, or *SUP99* plasmid were serially diluted and spotted onto a control YEPD plate and a YEPD plate containing 15 mM Caffeine and grown at 30°C for indicated days.

## DISCUSSION

Here we leverage the eukaryotic *S. cerevisiae* model system to unmask the mechanism(s) of action for known human oncohistone mutations. Budding yeast is an attractive model system for these experiments as histones are highly conserved, the budding yeast genome is streamlined with only two H3 genes instead of the 15 H3 genes present in the mammalian genome, and essential biological pathways that contribute to growth and metabolism are evolutionarily conserved. Because the *S. cerevisiae* genome contains only two copies of histone H3, *HHT1* and *HHT2*, this model enables robust genetic screening to: 1) identify pathways and biological processes that are altered to support oncogenicity in mammals; and 2) apply this information to develop and design rational therapeutics for the treatment of oncohistone-driven cancers.

Here we focus on the defined oncohistones H3K36M, H3G34W/L/R/V, and the histone missense mutation H3K36R. While not oncogenic, H3K36R serves as a model to explore how a conservative amino acid change at this position alters function (21). Initial experiments compare the growth of yeast cells that express these histone H3 variants as the sole copy of H3. The data reveal distinct growth phenotypes between H3K36 and H3G34 mutants (Figure 2), suggesting that while missense mutations that alter H3G34 reduce neighboring H3K36 methylation (Figure 1E), the functional dynamics are more complex. Furthermore, while H3K36 mutant cells show growth defects on plates with caffeine, they show no growth defect on plates containing hydroxyurea. In contrast, the H3G34 mutant cells all show growth defects in the presence of caffeine. Even cells with a significant decrease in the level of H3K36me3 such as H3G34V (∼25% of control H3K36me3) show no detectable growth defect on plates containing caffeine. These results align with recent studies that performed a similar analysis using *S. pombe* (17,18). The differences observed between oncohistone variants in yeast are also congruent with the different types of cancers linked to somatic mutations that alter H3K36 or H3G34 (2). Combined, these data suggest that additional analyses are required to uncover precisely how different amino acid changes within histones alter biological pathways to drive oncogenesis.

To define the biological pathways altered by different histone mutations, we exploited yeast genetics to perform a high copy suppressor screen. In this screen, we identified four candidate suppressors – the lysine acetyltransferase, Esa1, the gene expression regulator, Tos4, the m^6^A RNA binding protein Pho92, and the cyclin-dependent kinase, Sgv1/Bur1-that rescue the caffeine-associated growth defects that are exhibited by histone H3K36 missense mutations, which are known oncogenic drivers in cancer. The suppressors identified primarily suppress growth on caffeine with minimal effects on the other drugs that impair growth of the different histone models, suggesting that different biological pathways, which are impacted by the oncohistone missense mutations, can be defined with this genetic approach. Caffeine induces a stress response that depends on the Tor pathway (31), which is evolutionarily conserved as the mTOR pathway in humans (55). Identifying genes involved in suppressing growth defects associated with Tor pathway-induced stress is biologically relevant to human cancers, as the mammalian PI3K/AKT/mTOR signaling pathway mediates essential biological processes that are frequently deregulated in cancer: cell growth, survival, proliferation, and metabolism (56).

The high copy suppressor screen identified the histone H4/H2A lysine acetyltransferase Esa1 as a suppressor of caffeine sensitive growth of H3K36M and H3K36R histone mutant cells (Fig 2 & 3). The lysine acetyltransferase activity of Esa1 is required for the suppression as two independent catalytically inactive mutants of Esa1, Esa1 C340S and Esa1 E338Q (44,47), do not suppress the caffeine sensitive growth of H3K36R mutant cells. Interestingly, di- and trimethylation of histone H3K36 stimulates the interaction of the NuA4 lysine acetylation complex, which contains the Esa1 catalytic subunit, with nucleosomes and NuA4 acetylation of lysine residues in the histone H4 tail has also been observed to stimulate the interaction of the SAGA lysine acetylation complex with nucleosomes, enhancing acetylation of histone H3 (57). Overexpression of Esa1 in H3K36R mutant cells could therefore increase acetylation of histone H4 to enhance SAGA recruitment to nucleosomes to restore optimal acetylation of histone H3.

Crosstalk between H4 and/or H2A acetylation and H3K36 methylation could potentially impact cell growth or other pro-oncogenic properties that are abrogated in the absence of Esa1 enzymatic activity. However, the human homologue of Esa1, KAT5/TIP60, has additional non-histone substrates (58) that may impact biological function in high copy suppressor assays such as those employed here. TIP60 is involved in DNA repair through the acetylation of non-histone proteins such as the ATM kinase (59). Future experiments could include determining whether Esa1 suppresses growth in the presence of DNA damaging agents such as hydroxyurea, and whether this is dependent on Esa1 acetyltransferase activity. Ultimately, defining the relevant targets of Esa1 that confer growth suppression will lend insight into the biological pathways altered by oncogenic changes at H3K36.

The high copy screen also identified the gene expression regulator Tos4 as a potent suppressor of the caffeine sensitive growth of H3K36 mutant cells. Tos4 binds to the yeast histone deacetylase (HDAC) complexes Rpd3L and Set3 via a forkhead-associated (FHA) domain (Fig 4A) (39,40,60), which ultimately leads to histone deacetylation. However, amino acids substitutions that interfere with Rpd3L/Set3 complex binding to the FHA domain of Tos4 did not affect Tos4-mediated H3K36 mutant-growth suppression (Fig 4B), suggesting that Rpd3L/Set3-independent functions of Tos4 mediate suppression. This model is supported by the fact that we did not independently identify Rpd3L or Set3 complex genes as growth suppressors in our initial high copy suppressor screen. Pho92, an RNA “reader” protein that recognizes and binds m^6^A-containing RNA (48) is another suppressor identified in this screen. Amino acid substitutions within the reader domain that render Pho92 incapable of binding m^6^A RNA do not alter Pho92-mediated growth suppression of the H3K36R mutant growth, suggesting the mechanism(s) by which Pho92 mediates suppression is independent of recognizing and/or binding m^6^A RNA. Because Pho92 is not evolutionarily conserved between yeast and mammals, Pho92 suppression may be less likely to inform mechanism(s) of oncogenicity for H3K36-mutant tumors. Collectively, more research is needed to mechanistically define how some suppressors such as Tos4 and Pho92 suppress growth defects of H3K36-mutant cells.

The high copy suppressor screen also identified a clone containing nearly all of the *S. cerevisiae* kinase Sgv1/Bur1 open reading frame, a homologue of human CDK9 (Fig 5B). Previous studies have defined functional links between Sgv1/Bur1 and H3K36 methylation, demonstrating that a functional Sgv1/Bur1 protein is required for normal H3K36 methylation (61). While the slightly truncated *SGV1* genomic suppressor identified (*SUP3*) suppresses the caffeine sensitive growth of H3K36R mutant cells, the full-length *SGV1* clone does not. The genomic Sgv1 clone that was identified as a suppressor is truncated, lacking the two C-terminal amino acids (LY) and fused to an additional eight amino acids (PRVPSSNS) prior to reaching the next natural stop codon. As appending a C-terminal Myc tag with a very distinct sequence (EQKLISEEDL) to Sgv1 also suppresses the caffeine-sensitive growth of H3K36 mutant cells (Fig. 5B,C), the nature of the change to the C-terminus of Sgv1 is clearly not important. Furthermore, both the modified C-terminal end of the protein and the catalytic activity are required for suppression, suggesting a dominant function that relies on Sgv1 kinase activity. The C-terminal domain of *S. cerevisiae* Sgv1 is not evolutionarily conserved in mammalian CDK9 orthologs, but the C-terminal domain of Sgv1 is implicated in interaction with the RPA protein to ensure genome stability (62). In addition, the C-terminal domain of Sgv1 does share identity with the C-terminal domain of *S. pombe* Cdk9, which interacts with the mRNA capping/RNA 5’-triphophatase enzyme, Pct1 (Cet1 in *S. cerevisiae*) (63,64). Collectively, these data suggest that we may have identified a regulatory role for the C-terminal end of Sgv1 that depends on active kinase activity. Clearly further studies will be required to understand the interplay of Sgv1 and H3K36 oncohistone models.

The high copy suppressor screen performed here provides a streamlined and rapid approach to identify genetic vulnerabilities that may be therapeutically actionable. Two of the suppressors identified, Esa1 and Sgv1, encode enzymes that have been targeted in the corresponding pathways in human cells as potential therapies. Expression of KAT5/TIP60 lysine acetyl transferase, the human ortholog of Esa1, is deregulated in prostate and other cancers (65). The preclinical TIP60 acetyltransferase inhibitors Nu9056 and TH1834 reduce DNA-damage induced ATM phosphorylation (66). Thus, TIP60 inhibition may be a viable therapeutic strategy for cancers dependent on TIP60 enzymatic activity. This experimental approach in budding yeast provides the first evidence that H3K36-mutant tumors may be candidates for TIP60 inhibition, but subsequent experimentation in H3K36 mutant human cancer cell lines is required to test this hypothesis. The human homologue of Sgv1, CDK9 (51), binds to Cyclin T to form the positive transcription elongation factor complex (P-TEFb); CDK9 kinase activity is thus critical for RNA-pol II-directed transcription (67). CDK9 is deregulated in hematological malignancies and other cancer types, prompting the development of CDK9 inhibitors (68). However, understanding the mechanism(s) by which CDK9 inhibition contributes to anti-cancer effects is still under investigation. The CDK9 inhibitor AZD4573 is the first CDK9-selective inhibitor (69) and has entered clinical trials for hematological cancers. The anti-tumorigenic activity of CDK9 inhibition is thought to be due to the depletion of the anti-apoptotic protein MCL-1 (70). More recently, H3K27M oncohistone-expressing diffuse midline gliomas (DMGs) have been shown to deregulate the expression of AFF4, a scaffolding protein involved in transcriptional elongation, which CDK9/CyclinT regulates (71). These preclinical studies have prompted the clinical testing of AZD4573 in hematologic malignancies (72,73) and suggest that H3K27M-mutant tumors that are characterized by AFF4 upregulation are candidates for clinical CDK9 inhibition (73). While through a divergent mechanism, our data suggest that CDK9 inhibition may be beneficial for H3K36-mutant head and neck cancers and chondroblastomas, although further research is necessary to test this hypothesis.

Taken together, the results from this suppressor screen exploiting the budding yeast model both define novel cellular pathways that could be altered by missense mutations in histones that drive oncogenesis and uncover links to potential therapeutic avenues. Future studies will help to define the mechanisms by which the suppressors rescue the growth of the oncohistone yeast models, which may further define how these pathways could be targeted for therapeutic approaches.

## ACKNOWLEDGEMENTS

We thank members of the Corbett lab for project discussions and experimental insight. This study was supported by grants from the NIH (R21 CA256456 to A.H.C. and J.M.S.), a Post-doctoral Enrichment Program (PDEP) Award from Burroughs Wellcome to L.D.L. L.D.L was supported by the NIH IRACDA program (K12 GM000680). H.N. and R.A. participated in the Emory NIH Initiative for Maximizing Student Development (IMSD) Program (GM125598). V.G. was supported by the Laney Graduate School-Summer Opportunity for Academic Research (SOAR) Program.

## References

1. Strahl, B. D., and Allis, C. D. (2000) The language of covalent histone modifications. Nature 403, 41–45

2. Wan, Y. C. E., Liu, J., and Chan, K. M. (2018) Histone H3 Mutations in Cancer. Curr Pharmacol Rep 4, 292–300

3. Harutyunyan, A. S., Krug, B., Chen, H., Papillon-Cavanagh, S., Zeinieh, M., De Jay, N., Deshmukh, S., Chen, C. C. L., Belle, J., Mikael, L. G., Marchione, D. M., Li, R., Nikbakht, H., Hu, B., Cagnone, G., Cheung, W. A., Mohammadnia, A., Bechet, D., Faury, D., McConechy, M. K., Pathania, M., Jain, S. U., Ellezam, B., Weil, A. G., Montpetit, A., Salomoni, P., Pastinen, T., Lu, C., Lewis, P. W., Garcia, B. A., Kleinman, C. L., Jabado, N., and Majewski, J. (2019) H3K27M induces defective chromatin spread of PRC2-mediated repressive H3K27me2/me3 and is essential for glioma tumorigenesis. Nat Commun 10, 1262

4. Justin, N., Zhang, Y., Tarricone, C., Martin, S. R., Chen, S., Underwood, E., De Marco, V., Haire, L. F., Walker, P. A., Reinberg, D., Wilson, J. R., and Gamblin, S. J. (2016) Structural basis of oncogenic histone H3K27M inhibition of human polycomb repressive complex 2. Nat Commun 7, 11316

5. Lewis, P. W., Muller, M. M., Koletsky, M. S., Cordero, F., Lin, S., Banaszynski, L. A., Garcia, B. A., Muir, T. W., Becher, O. J., and Allis, C. D. (2013) Inhibition of PRC2 activity by a gain-of-function H3 mutation found in pediatric glioblastoma. Science 340, 857–861

6. Lu, C., Jain, S. U., Hoelper, D., Bechet, D., Molden, R. C., Ran, L., Murphy, D., Venneti, S., Hameed, M., Pawel, B. R., Wunder, J. S., Dickson, B. C., Lundgren, S. M., Jani, K. S., De Jay, N., Papillon-Cavanagh, S., Andrulis, I. L., Sawyer, S. L., Grynspan, D., Turcotte, R. E., Nadaf, J., Fahiminiyah, S., Muir, T. W., Majewski, J., Thompson, C. B., Chi, P., Garcia, B. A., Allis, C. D., Jabado, N., and Lewis, P. W. (2016) Histone H3K36 mutations promote sarcomagenesis through altered histone methylation landscape. Science 352, 844–849

7. Yang, S., Zheng, X., Lu, C., Li, G. M., Allis, C. D., and Li, H. (2016) Molecular basis for oncohistone H3 recognition by SETD2 methyltransferase. Genes Dev 30, 1611–1616

8. Braastad, C. D., Hovhannisyan, H., van Wijnen, A. J., Stein, J. L., and Stein, G. S. (2004) Functional characterization of a human histone gene cluster duplication. Gene 342, 35–40

9. Nacev, B. A., Feng, L., Bagert, J. D., Lemiesz, A. E., Gao, J., Soshnev, A. A., Kundra, R., Schultz, N., Muir, T. W., and Allis, C. D. (2019) The expanding landscape of ‘oncohistone’ mutations in human cancers. Nature 567, 473–478

10. Taylor, E. L., and Westendorf, J. J. (2021) Histone Mutations and Bone Cancers. Adv. Exp. Med. Biol. 1283, 53–62

11. Zhang, Y., Zhou, L., Safran, H., Borsuk, R., Lulla, R., Tapinos, N., Seyhan, A. A., and El-Deiry, W. S. (2021) EZH2i EPZ-6438 and HDACi vorinostat synergize with ONC201/TIC10 to activate integrated stress response, DR5, reduce H3K27 methylation, ClpX and promote apoptosis of multiple tumor types including DIPG. Neoplasia 23, 792-810

12. Yusufova, N., Kloetgen, A., Teater, M., Osunsade, A., Camarillo, J. M., Chin, C. R., Doane, A. S., Venters, B. J., Portillo-Ledesma, S., Conway, J., Phillip, J. M., Elemento, O., Scott, D. W., Béguelin, W., Licht, J. D., Kelleher, N. L., Staudt, L. M., Skoultchi, A. I., Keogh, M. C., Apostolou, E., Mason, C. E., Imielinski, M., Schlick, T., David, Y., Tsirigos, A., Allis, C. D., Soshnev, A. A., Cesarman, E., and Melnick, A. M. (2021) Histone H1 loss drives lymphoma by disrupting 3D chromatin architecture. Nature 589, 299–305

13. Wan, Y. C. E., and Chan, K. M. (2021) Histone H2B Mutations in Cancer. Biomedicines 9

14. Kang, T. Z. E., Zhu, L., Yang, D., Ding, D., Zhu, X., Wan, Y. C. E., Liu, J., Ramakrishnan, S., Chan, L. L., Chan, S. Y., Wang, X., Gan, H., Han, J., Ishibashi, T., Li, Q., and Chan, K. M. (2021) The elevated transcription of ADAM19 by the oncohistone H2BE76K contributes to oncogenic properties in breast cancer. J Biol Chem 296, 100374

15. Duina, A. A., Miller, M. E., and Keeney, J. B. (2014) Budding yeast for budding geneticists: a primer on the Saccharomyces cerevisiae model system. Genetics 197, 33–48

16. Bagert, J. D., Mitchener, M. M., Patriotis, A. L., Dul, B. E., Wojcik, F., Nacev, B. A., Feng, L., Allis, C. D., and Muir, T. W. (2021) Oncohistone mutations enhance chromatin remodeling and alter cell fates. Nat Chem Biol 17, 403–411

17. Lowe, B. R., Yadav, R. K., Henry, R. A., Schreiner, P., Matsuda, A., Fernandez, A. G., Finkelstein, D., Campbell, M., Kallappagoudar, S., Jablonowski, C. M., Andrews, A. J., Hiraoka, Y., and Partridge, J. F. (2021) Surprising phenotypic diversity of cancer-associated mutations of Gly 34 in the histone H3 tail. Elife 10

18. Yadav, R. K., Jablonowski, C. M., Fernandez, A. G., Lowe, B. R., Henry, R. A., Finkelstein, D., Barnum, K. J., Pidoux, A. L., Kuo, Y. M., Huang, J., O’Connell, M. J., Andrews, A. J., Onar-Thomas, A., Allshire, R. C., and Partridge, J. F. (2017) Histone H3G34R mutation causes replication stress, homologous recombination defects and genomic instability in S. pombe. Elife 6

19. Hyland, E. M., Cosgrove, M. S., Molina, H., Wang, D., Pandey, A., Cottee, R. J., and Boeke, J. D. (2005) Insights into the role of histone H3 and histone H4 core modifiable residues in Saccharomyces cerevisiae. Molecular and cellular biology 25, 10060–10070

20. Meers, M. P., Henriques, T., Lavender, C. A., McKay, D. J., Strahl, B. D., Duronio, R. J., Adelman, K., and Matera, A. G. (2017) Histone gene replacement reveals a post-transcriptional role for H3K36 in maintaining metazoan transcriptome fidelity. Elife 6

21. McDaniel, S. L., Hepperla, A. J., Huang, J., Dronamraju, R., Adams, A. T., Kulkarni, V. G., Davis, I. J., and Strahl, B. D. (2017) H3K36 Methylation Regulates Nutrient Stress Response in Saccharomyces cerevisiae by Enforcing Transcriptional Fidelity. Cell Rep 19, 2371–2382

22. Sorenson, M. R., Jha, D. K., Ucles, S. A., Flood, D. M., Strahl, B. D., Stevens, S. W., and Kress, T. L. (2016) Histone H3K36 methylation regulates pre-mRNA splicing in Saccharomyces cerevisiae. RNA Biol. 13, 412–426

23. Kaczmarek Michaels, K., Mohd Mostafa, S., Ruiz Capella, J., and Moore, Claire L. (2020) Regulation of alternative polyadenylation in the yeast Saccharomyces cerevisiae by histone H3K4 and H3K36 methyltransferases. Nucleic Acids Res. 48, 5407–5425

24. Venkatesh, S., Li, H., Gogol, M. M., and Workman, J. L. (2016) Selective suppression of antisense transcription by Set2-mediated H3K36 methylation. Nature Communications 7, 13610

25. Adams, A., Gottschling, D. E., Kaiser, C. A., and Stearns, T. (1997) Methods in Yeast Genetics, Cold Spring Harbor Laboratory Press, Cold Spring Harbor

26. Sambrook, J., Fritsch, E. F., and Maniatis, T. (1989) Molecular Cloning: A Laboratory Manual, Second ed., Cold Spring Harbor Laboratory Press, Cold Spring Harbor, New York

27. Duina, A. A., and Turkal, C. E. (2017) Targeted in Situ Mutagenesis of Histone Genes in Budding Yeast. J Vis Exp

28. Johnson, P., Mitchell, V., McClure, K., Kellems, M., Marshall, S., Allison, M. K., Lindley, H., Nguyen, H. T., Tackett, J. E., and Duina, A. A. (2015) A systematic mutational analysis of a histone H3 residue in budding yeast provides insights into chromatin dynamics. G3 (Bethesda) 5, 741-749

29. Zhang, C. J., Cavenagh, M. M., and Kahn, R. A. (1998) A family of Arf effectors defined as suppressors of the loss of Arf function in the yeast Saccharomyces cerevisiae. J Biol Chem 273, 19792–19796

30. Hill, J. E., Myers, A. M., Koerner, T. J., and Tzagoloff, A. (1986) Yeast/E. coli shuttle vectors with multiple unique restriction sites. Yeast 2, 163–167

31. Kuranda, K., Leberre, V., Sokol, S., Palamarczyk, G., and François, J. (2006) Investigating the caffeine effects in the yeast Saccharomyces cerevisiae brings new insights into the connection between TOR, PKC and Ras/cAMP signalling pathways. Molecular microbiology 61, 1147–1166

32. Hoyos-Manchado, R., Reyes-Martín, F., Rallis, C., Gamero-Estévez, E., Rodríguez-Gómez, P., Quintero-Blanco, J., Bähler, J., Jiménez, J., and Tallada, V. A. (2017) RNA metabolism is the primary target of formamide in vivo. Sci Rep 7, 15895

33. Slater, M. L. (1973) Effect of reversible inhibition of deoxyribonucleic acid synthesis on the yeast cell cycle. J Bacteriol 113, 263–270

34. Strahl, B. D., Grant, P. A., Briggs, S. D., Sun, Z. W., Bone, J. R., Caldwell, J. A., Mollah, S., Cook, R. G., Shabanowitz, J., Hunt, D. F., and Allis, C. D. (2002) Set2 is a nucleosomal histone H3-selective methyltransferase that mediates transcriptional repression. Molecular and cellular biology 22, 1298–1306

35. Irie, K., Nomoto, S., Miyajima, I., and Matsumoto, K. (1991) SGV1 encodes a CDC28/cdc2-related kinase required for a G alpha subunit-mediated adaptive response to pheromone in S. cerevisiae. Cell 65, 785–795

36. Prelich, G., and Winston, F. (1993) Mutations that suppress the deletion of an upstream activating sequence in yeast: involvement of a protein kinase and histone H3 in repressing transcription in vivo. Genetics 135, 665–676

37. Clarke, A. S., Lowell, J. E., Jacobson, S. J., and Pillus, L. (1999) Esa1p is an essential histone acetyltransferase required for cell cycle progression. Molecular and cellular biology 19, 2515–2526

38. Horak, C. E., Luscombe, N. M., Qian, J., Bertone, P., Piccirrillo, S., Gerstein, M., and Snyder, M. (2002) Complex transcriptional circuitry at the G1/S transition in Saccharomyces cerevisiae. Genes Dev 16, 3017–3033

39. Cooke, S. L., Soares, B. L., Müller, C. A., Nieduszynski, C. A., Bastos de Oliveira, F. M., and de Bruin, R. A. M. (2021) Tos4 mediates gene expression homeostasis through interaction with HDAC complexes independently of H3K56 acetylation. J Biol Chem 296, 100533

40. Bastos de Oliveira, F. M., Harris, M. R., Brazauskas, P., de Bruin, R. A., and Smolka, M. B. (2012) Linking DNA replication checkpoint to MBF cell-cycle transcription reveals a distinct class of G1/S genes. The EMBO journal 31, 1798–1810

41. Kang, H. J., Jeong, S. J., Kim, K. N., Baek, I. J., Chang, M., Kang, C. M., Park, Y. S., and Yun, C. W. (2014) A novel protein, Pho92, has a conserved YTH domain and regulates phosphate metabolism by decreasing the mRNA stability of PHO4 in Saccharomyces cerevisiae. Biochem J 457, 391-400

42. Smith, E. R., Eisen, A., Gu, W., Sattah, M., Pannuti, A., Zhou, J., Cook, R. G., Lucchesi, J. C., and Allis, C. D. (1998) ESA1 is a histone acetyltransferase that is essential for growth in yeast. Proc Natl Acad Sci U S A 95, 3561–3565

43. Doyon, Y., Selleck, W., Lane, W. S., Tan, S., and Côté, J. (2004) Structural and functional conservation of the NuA4 histone acetyltransferase complex from yeast to humans. Molecular and cellular biology 24, 1884–1896

44. Yan, Y., Barlev, N. A., Haley, R. H., Berger, S. L., and Marmorstein, R. (2000) Crystal structure of yeast Esa1 suggests a unified mechanism for catalysis and substrate binding by histone acetyltransferases. Mol. Cell 6, 1195–1205

45. Lafon, A., Chang, C. S., Scott, E. M., Jacobson, S. J., and Pillus, L. (2007) MYST opportunities for growth control: yeast genes illuminate human cancer gene functions. Oncogene 26, 5373–5384

46. Eissenberg, J. C. (2012) Structural biology of the chromodomain: form and function. Gene 496, 69–78

47. Decker, P. V., Yu, D. Y., Iizuka, M., Qiu, Q., and Smith, M. M. (2008) Catalytic-site mutations in the MYST family histone Acetyltransferase Esa1. Genetics 178, 1209–1220

48. Shi, R., Ying, S., Li, Y., Zhu, L., Wang, X., and Jin, H. (2021) Linking the YTH domain to cancer: the importance of YTH family proteins in epigenetics. Cell Death Dis. 12, 346

49. Xu, C., Liu, K., Ahmed, H., Loppnau, P., Schapira, M., and Min, J. (2015) Structural Basis for the Discriminative Recognition of N6-Methyladenosine RNA by the Human YT521-B Homology Domain Family of Proteins. J Biol Chem 290, 24902–24913

50. Theler, D., Dominguez, C., Blatter, M., Boudet, J., and Allain, F. H.-T. (2014) Solution structure of the YTH domain in complex with N6-methyladenosine RNA: a reader of methylated RNA. Nucleic Acids Res. 42, 13911–13919

51. Chun, Y., Joo, Y. J., Suh, H., Batot, G., Hill, C. P., Formosa, T., and Buratowski, S. (2019) Selective Kinase Inhibition Shows That Bur1 (Cdk9) Phosphorylates the Rpb1 Linker In Vivo. Molecular and cellular biology 39

52. Malumbres, M. (2014) Cyclin-dependent kinases. Genome Biol. 15, 122

53. Keogh, M. C., Podolny, V., and Buratowski, S. (2003) Bur1 kinase is required for efficient transcription elongation by RNA polymerase II. Molecular and cellular biology 23, 7005–7018

54. Zhou, K., Kuo, W. H., Fillingham, J., and Greenblatt, J. F. (2009) Control of transcriptional elongation and cotranscriptional histone modification by the yeast BUR kinase substrate Spt5. Proc Natl Acad Sci U S A 106, 6956–6961

55. Tatebe, H., and Shiozaki, K. (2017) Evolutionary Conservation of the Components in the TOR Signaling Pathways. Biomolecules 7

56. Saxton, R. A., and Sabatini, D. M. (2017) mTOR Signaling in Growth, Metabolism, and Disease. Cell 168, 960–976

57. Ginsburg, D. S., Anlembom, T. E., Wang, J., Patel, S. R., Li, B., and Hinnebusch, A. G. (2014) NuA4 links methylation of histone H3 lysines 4 and 36 to acetylation of histones H4 and H3. J Biol Chem 289, 32656–32670

58. Sapountzi, V., and Côté, J. (2011) MYST-family histone acetyltransferases: beyond chromatin. Cell. Mol. Life Sci. 68, 1147–1156

59. Sun, Y., Xu, Y., Roy, K., and Price, B. D. (2007) DNA damage-induced acetylation of lysine 3016 of ATM activates ATM kinase activity. Molecular and cellular biology 27, 8502–8509

60. Hofmann, K., and Bucher, P. (1995) The FHA domain: a putative nuclear signalling domain found in protein kinases and transcription factors. Trends Biochem. Sci. 20, 347–349

61. Chu, Y., Sutton, A., Sternglanz, R., and Prelich, G. (2006) The BUR1 cyclin-dependent protein kinase is required for the normal pattern of histone methylation by SET2. Molecular and cellular biology 26, 3029–3038

62. Clausing, E., Mayer, A., Chanarat, S., Müller, B., Germann, S. M., Cramer, P., Lisby, M., and Strässer, K. (2010) The transcription elongation factor Bur1-Bur2 interacts with replication protein A and maintains genome stability during replication stress. J Biol Chem 285, 41665–41674

63. Pei, Y., and Shuman, S. (2003) Characterization of the Schizosaccharomyces pombe Cdk9/Pch1 protein kinase: Spt5 phosphorylation, autophosphorylation, and mutational analysis. J Biol Chem 278, 43346–43356

64. Takagi, T., Cho, E. J., Janoo, R. T., Polodny, V., Takase, Y., Keogh, M. C., Woo, S. A., Fresco-Cohen, L. D., Hoffman, C. S., and Buratowski, S. (2002) Divergent subunit interactions among fungal mRNA 5’-capping machineries. Eukaryot Cell 1, 448–457

65. Shiota, M., Yokomizo, A., Masubuchi, D., Tada, Y., Inokuchi, J., Eto, M., Uchiumi, T., Fujimoto, N., and Naito, S. (2010) Tip60 promotes prostate cancer cell proliferation by translocation of androgen receptor into the nucleus. The Prostate 70, 540–554

66. Gao, C., Bourke, E., Scobie, M., Famme, M. A., Koolmeister, T., Helleday, T., Eriksson, L. A., Lowndes, N. F., and Brown, J. A. L. (2014) Rational design and validation of a Tip60 histone acetyltransferase inhibitor. Scientific Reports 4, 5372

67. Bacon, C. W., and D’Orso, I. (2019) CDK9: a signaling hub for transcriptional control. Transcription 10, 57–75

68. Mandal, R., Becker, S., and Strebhardt, K. (2021) Targeting CDK9 for Anti-Cancer Therapeutics. Cancers (Basel) 13, 2181

69. Cidado, J., Boiko, S., Proia, T., Ferguson, D., Criscione, S. W., San Martin, M., Pop-Damkov, P., Su, N., Roamio Franklin, V. N., Sekhar Reddy Chilamakuri, C., D’Santos, C. S., Shao, W., Saeh, J. C., Koch, R., Weinstock, D. M., Zinda, M., Fawell, S. E., and Drew, L. (2020) AZD4573 Is a Highly Selective CDK9 Inhibitor That Suppresses MCL-1 and Induces Apoptosis in Hematologic Cancer Cells. Clin. Cancer Res. 26, 922–934

70. Dey, J., Deckwerth, T. L., Kerwin, W. S., Casalini, J. R., Merrell, A. J., Grenley, M. O., Burns, C., Ditzler, S. H., Dixon, C. P., Beirne, E., Gillespie, K. C., Kleinman, E. F., and Klinghoffer, R. A. (2017) Voruciclib, a clinical stage oral CDK9 inhibitor, represses MCL-1 and sensitizes high-risk Diffuse Large B-cell Lymphoma to BCL2 inhibition. Sci Rep 7, 18007

71. Dahl, N. A., Danis, E., Balakrishnan, I., Wang, D., Pierce, A., Walker, F. M., Gilani, A., Serkova, N. J., Madhavan, K., Fosmire, S., Green, A. L., Foreman, N. K., Venkataraman, S., and Vibhakar, R. (2020) Super Elongation Complex as a Targetable Dependency in Diffuse Midline Glioma. Cell Rep 31, 107485

72. Rule, S., Kater, A. P., Brümmendorf, T. H., Fegan, C., Kaiser, M., Radford, J. A., Stilgenbauer, S., Kayser, S., Dyer, M. J., Brossart, P., Cidado, J., Drew, L., Birkett, J., Davies, A., Shao, W., and Brugger, W. (2018) A phase 1, open-label, multicenter, non-randomized study to assess the safety, tolerability, pharmacokinetics, and preliminary antitumor activity of AZD4573, a potent and selective CDK9 inhibitor, in subjects with relapsed or refractory hematological malignancies. J. Clin. Oncol. 36, TPS7588-TPS7588

73. Barlaam, B., Casella, R., Cidado, J., Cook, C., De Savi, C., Dishington, A., Donald, C. S., Drew, L., Ferguson, A. D., Ferguson, D., Glossop, S., Grebe, T., Gu, C., Hande, S., Hawkins, J., Hird, A. W., Holmes, J., Horstick, J., Jiang, Y., Lamb, M. L., McGuire, T. M., Moore, J. E., O’Connell, N., Pike, A., Pike, K. G., Proia, T., Roberts, B., San Martin, M., Sarkar, U., Shao, W., Stead, D., Sumner, N., Thakur, K., Vasbinder, M. M., Varnes, J. G., Wang, J., Wang, L., Wu, D., Wu, L., Yang, B., and Yao, T. (2020) Discovery of AZD4573, a Potent and Selective Inhibitor of CDK9 That Enables Short Duration of Target Engagement for the Treatment of Hematological Malignancies. J. Med. Chem. 63, 15564–15590

