## Supplemental Table 1 for "A Budding Yeast Model and Screen to Define the Functional Consequences of Oncogenic Histone Missense Mutations"

**Supplemental Table 1** Lemon *et al.*

| Strain/Plasmid | Description | Source |
|---|---|---|
| Wildtype (yAAD1253) | *MATα, ura3-52, leu2Δ1, his3Δ200, lys2-128δ* | (1) |
| *hht2*Δ (yAAD165) | *MATa, ura3-52, leu2Δ1, his3Δ200, trp1Δ63, lys2-128δ, hht2Δ::URA3:TRP1* | (1) |
| *hht1*Δ (ACY2818) | *MATα, ura3-52, leu2Δ1 his3Δ200, lys2-128δ, hht1Δ::kanMX* | This Study |
| *hht2-K36R* (ACY2816) | *MATa, ura3-52, leu2Δ1, his3Δ200, trp1Δ63, lys2-128δ, hht2-K36R* | This Study |
| *hht2-K36R hht1*Δ (ACY2821) | *MATa, ura3-52, leu2Δ1, his3Δ200, trp1Δ63, lys2-128δ, hht2-K36R, hht1Δ::kanMX* | This Study |
| *hht2-K36M* (ACY2830) | *MATa, ura3-52, leu2Δ1, his3Δ200, trp1Δ63, lys2-128δ, hht2-K36M* | This Study |
| *hht2-K36M hht1*Δ (ACY2822) | *MATa, ura3-52, leu2Δ1, his3Δ200, trp1Δ63, lys2-128δ, hht2-K36M, hht1Δ::kanMX* | This Study |
| *hht2-G34W* (ACY2823) | *MATa, ura3-52, leu2Δ1, his3Δ200, trp1Δ63, lys2-128δ, hht2-G34W* | This Study |
| *hht2-G34W hht1*Δ (ACY2825) | *MATa, ura3-52, leu2Δ1, his3Δ200, trp1Δ63, lys2-128δ, hht2-G34W, hht1Δ::kanMX* | This Study |
| *hht2-G34L* (ACY2831) | *MATa, ura3-52, leu2Δ1, his3Δ200, trp1Δ63, lys2-128δ, hht2-G34L* | This Study |
| *hht2-G34L hht1*Δ (ACY2833) | *MATa, ura3-52, leu2Δ1, his3Δ200, trp1Δ63, lys2-128δ, hht2-G34L, hht1Δ::kanMX* | This Study |
| *hht2-G34R* (ACY2838) | *MATa, ura3-52, leu2Δ1, his3Δ200, trp1Δ63, lys2-128δ, hht2-G34R* | This Study |
| *hht2-G34R hht1*Δ (ACY2840) | *MATa, ura3-52, leu2Δ1, his3Δ200, trp1Δ63, lys2-128δ, hht2-G34R, hht1Δ::kanMX* | This Study |
| *hht2-G34V* (ACY2841) | *MATa, ura3-52, leu2Δ1, his3Δ200, trp1Δ63, lys2-128δ, hht2-G34V* | This Study |
| *hht2-G34V hht1*Δ (ACY2846) | *MATa, ura3-52, leu2Δ1, his3Δ200, trp1Δ63, lys2-128δ, hht2-G34V, hht1Δ::kanMX* | This Study |
| *set2*Δ (ACY2851) | *MATα, ura3-52, leu2Δ1, his3Δ200, lys2-128δ, set2Δ::kanMX* | This Study |
| YEp352 (pAC29) | *URA3, 2μ, amp*$^R$ | (2) |
| *SUP3* (pAC4132) | *SGV1, URA3, 2μ, amp*$^R$ | This Study |
| *SUP54* (pAC4145) | *HHT2, HHF2, URA3, 2μ, amp*$^R$ | This Study |
| *SUP67* (pAC4149) | *ESA1, URA3, 2μ, amp*$^R$ | This Study |
| *SUP68* (pAC4150) | *TOS4, YLR184W, URA3, 2μ, amp*$^R$ | This Study |
| *SUP99* (pAC4160) | *PHO92, WIP1, BCS1, URA3, 2μ, amp*$^R$ | This Study |
| *HHF2* (pAC4199) | *HHF2, URA3, 2μ, amp*$^R$ | This Study |
| *HHT2* (pAC4201) | *HHT2, URA3, 2μ, amp*$^R$ | This Study |
| *HHT1* (pAC4200) | *HHT1, URA3, 2μ, amp*$^R$ | This Study |
| *ESA1* (pAC4190) | *ESA1, URA3, 2μ, amp*$^R$ | This Study |
| *esa1-C304S* (pAC4191) | *esa1-C304S, URA3, 2μ, amp*$^R$ | This Study |
| *esa1-E338Q* (pAC4192) | *esa1-E338Q, URA3, 2μ, amp*$^R$ | This Study |
| *TOS4* (pAC4196) | *TOS4, URA3, 2μ, amp*$^R$ | This Study |
| *tos4-R122A-N161A* (pAC4205) | *tos4-R122A-N161A, URA3, 2μ, amp*$^R$ | This Study |
| *PHO92* (pAC4193) | *PHO92, URA3, 2μ, amp*$^R$ | This Study |
| *pho92-W177A* (pAC4194) | *pho92-W177A, URA3, 2μ, amp*$^R$ | This Study |
| *pho92-W231A* (pAC4195) | *pho92-W231A, URA3, 2μ, amp*$^R$ | This Study |
| *SGV1* (pAC4187) | *SGV1, URA3, 2μ, amp*$^R$ | This Study |
| *sgv1-E107Q* (pAC4188) | *sgv1-E107Q, URA3; 2μ, amp*$^R$ | This Study |
| *sgv1-D213A* (pAC4189) | *sgv1-D213A, URA3; 2μ, amp*$^R$ | This Study |
| *sgv1-Δ2+8aa* (pAC4208) | sgv1-Δ2+8aa, URA3, 2μ, *amp*$^R$ | This Study |
| *SGV1-Myc* (pAC4209) | *SGV1-Myc, URA3, 2μ, amp*$^R$ | This Study |
| *sgv1-E107Q-Myc* (pAC4210) | *sgv1-E107Q-Myc, URA3, 2μ, amp*$^R$ | This Study |
| *sgv1-D213A-Myc* (pAC4211) | *sgv1-D213A-Myc, URA3, 2μ, amp*$^R$ | This Study |
| *SUP3-E107Q* (pAC4212) | *SUP3-E107Q, URA3, 2μ, amp*$^R$ | This Study |
| *SUP3-D213A* (pAC4213) | *SUP3-D213A, URA3, 2μ, amp*$^R$ | This Study |

Supplemental Figure 1

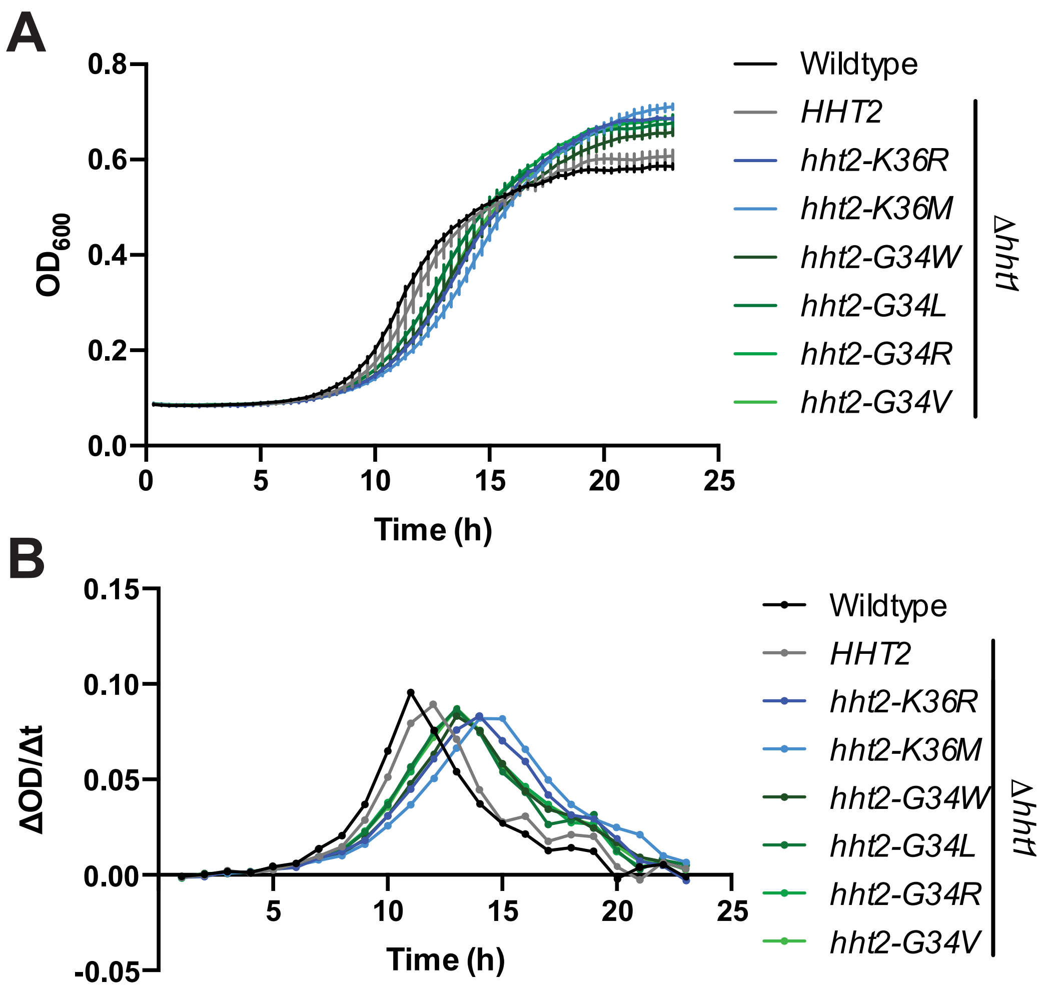


**Figure S1.** (A) Control Wildtype and *HHT2 hht1*Δ cells (black), H3K36 mutant cells (*hht2-K36R/M hht1*Δ) (blue), or H3G34 mutant cells (*hht2-G34W/L/R/V hht1*Δ) (green) were grown in YEPD liquid media and growth was assessed my measuring OD600 every 20 minutes in a plate reader for 24 hours. (B) The area under the growth curve was obtained for each of the mutants analyzed, revealing that the histone mutant cells achieve a higher biomass than either control.

### Supplemental Figure 2

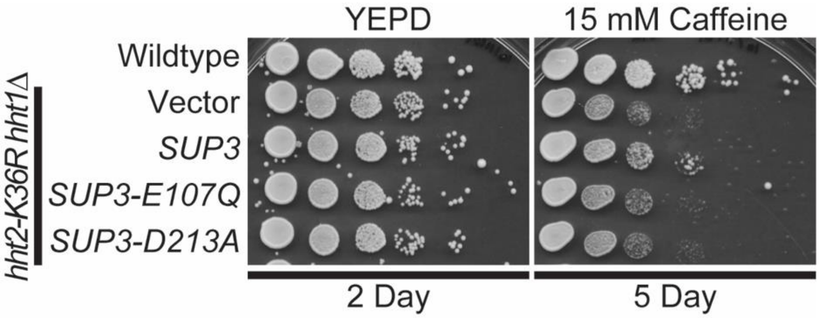


**Figure S2.** Both the catalytic activity of Sgv1 and a C-terminal extension are required for Sgv1-mediated suppression of H3K36R caffeine-sensitive growth. Using the original suppressor clone, termed *SUP3*, either of two Sgv1 amino acid substitutions that impair Sgv1 catalytic activity (E107Q or D213A) (3) engineered into the original suppressor clone, *SUP3*, identified abrogate suppression.

### Supplemental References


1. Duina, A. A., and Turkal, C. E. (2017) Targeted in Situ Mutagenesis of Histone Genes in Budding Yeast. *J Vis Exp*, 55263
2. Hill, J. E., Myers, A. M., Koerner, T. J., and Tzagoloff, A. (1986) Yeast/E. coli shuttle vectors with multiple unique restriction sites. *Yeast* **2**, 163-167
3. Keogh, M. C., Podolny, V., and Buratowski, S. (2003) Bur1 kinase is required for efficient transcription elongation by RNA polymerase II. *Molecular and cellular biology* **23**, 7005-7018
